# Koina: Democratizing machine learning for proteomics research

**DOI:** 10.1101/2024.06.01.596953

**Authors:** Ludwig Lautenbacher, Kevin L. Yang, Tobias Kockmann, Christian Panse, Matthew Chambers, Elias Kahl, Fengchao Yu, Wassim Gabriel, Dulguun Bold, Tobias Schmidt, Kai Li, Brendan MacLean, Alexey I. Nesvizhskii, Mathias Wilhelm

## Abstract

Recent developments in machine-learning (ML) and deep-learning (DL) have immense potential for applications in proteomics, such as generating spectral libraries, improving peptide identification, and optimizing targeted acquisition modes. Although new ML/DL models for various applications and peptide properties are frequently published, the rate at which these models are adopted by the community is slow, which is mostly due to technical challenges. We believe that, for the community to make better use of state-of-the-art models, more attention should be spent on making models easy to use and accessible by the community. To facilitate this, we developed Koina, an open-source containerized, decentralized and online-accessible high-performance prediction service that enables ML/DL model usage in any pipeline. Using the widely used FragPipe computational platform as example, we show how Koina can be easily integrated with existing proteomics software tools and how these integrations improve data analysis.

## Introduction

Recent developments in machine-learning (ML) and deep-learning (DL) have immense potential for applications in proteomics, such as generating optimized spectral libraries^1^ and improving peptide identification in DDA^2–6^, DIA^7^, and targeted acquisition modes^8^. Although new ML/DL models for various applications and peptide properties are frequently published, the community adopts only a few of them. This is largely due to a lack of findability, accessibility, interoperability, and reusability (FAIR)^9,10^ of most published machine-learning models. With limited model exchange formats and language-specific ML/DL frameworks mostly available in Python, access from other programming languages commonly used in data analysis like R, Java, JavaScript, and C# is very difficult. Furthermore, ML models, and especially DL models, commonly require access to specialized hardware, the requirements and price of which have steadily increased over the last years. This unintentionally divides the scientific community into those that can invest time and other resources to set up and maintain the necessary hardware and those that cannot. So far, solving this issue relied mostly on ML developers going out of their way to enable accessibility by developing custom solutions^6,11,12^. This comes with the common negative side effects of custom solutions, such as the reimplementation of already existing functionality, users having to adapt to new interfaces for every new model, and functionality that exists in one service but not another. Since these issues are common in all fields that apply ML/DL models, attempts have already been made to address them. Most commonly, this is done by developing a model repository containing pre-trained machine-learning models ready to be downloaded and applied directly^13–15^.

Here, we describe the initial release of Koina, a web-accessible model repository that facilitates access to high-performance machine-learning models predicting peptide properties used in computational proteomics analysis. We describe the design decisions and how they benefit ML/DL developers, downstream tool developers, and end users of said software tools. Koina goes one step beyond the classic model repository approach by providing models not only for offline use but making them available via web traffic. This combines the benefits of a model repository with the usability of a web service. We believe this approach holds significant potential to “democratize” machine-learning by enabling laboratories with limited access to high-performance computing to benefit from ML models in their data analysis as well. Koina is already integrated in the popular proteomics data analysis software packages FragPipe^4^, Skyline^16^, and Oktoberfest^2^. Furthermore, we show complementary Python and R packages that further simplify accessing Koina. Finally, we benchmark many of the Koina models in the FragPipe software suite with MSBooster^4^ for peptide-spectrum-match (PSM) rescoring and, anticipating a growing list of available models, provide a practical solution for finding optimal models for PSM rescoring.

## Results & Discussion

### Koina is a platform to democratize access to ML models for proteomics

Koina is a model repository enabling the remote execution of models. Predictions are generated as a response to HTTP/S requests, the standard protocol used for nearly all web traffic. As such, HTTP/S requests can be easily generated in any programming language without requiring specialized hardware. This design also enables users to share centralized hardware to utilize it more efficiently. It also allows for easy horizontal scaling depending on the demand of the user base (Fig 1a). To minimize the barrier of entry and “democratize” access to ML models, we provide a public network of Koina instances at koina.wilhelmlab.org. The computational workload is automatically distributed to processing nodes hosted at different research institutions and spin-offs across Europe. Each processing node provides computational resources to the service network, always aiming at just-in-time results delivery. In the spirit of open and collaborative science, we envision that this public Koina-Network can be scaled to meet the community’s needs by various research groups or institutions dedicating hardware. This can also vastly improve latency if servers are available geographically nearby. Alternatively, if data security is a concern, private instances within a local network can be easily deployed using the provided docker image (Fig 1a).

**Figure 1:**
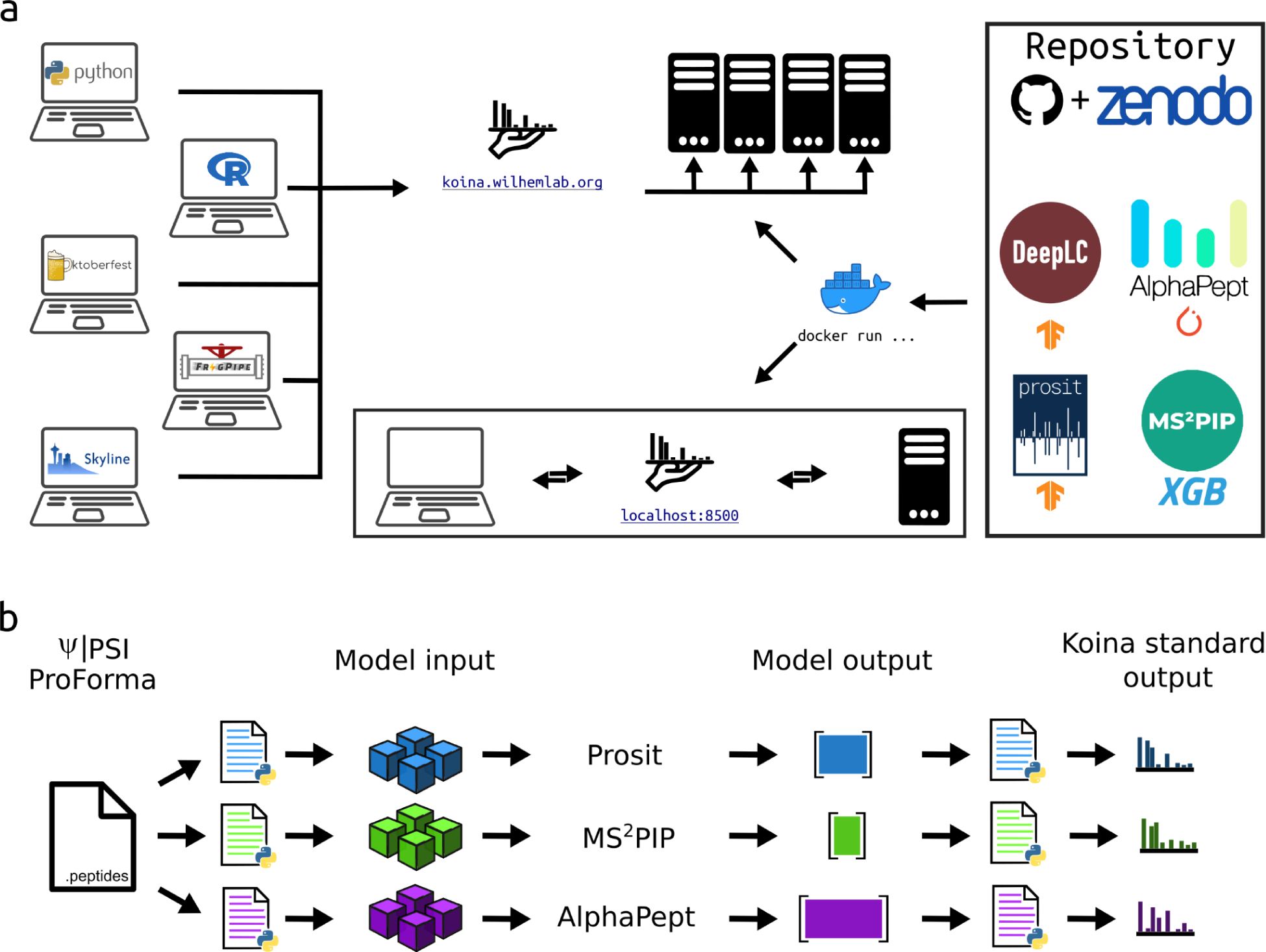
(a) Overview of the Koina web-accessible model repository. All code used for Koina is publicly available on GitHub. Model weights are stored on Zenodo and fetched dynamically on server startup. The provided docker image allows for easy scaling & deployment of private instances. The web service design of Koina allows requests from any source, such as KoinaPy (Python), KoinaR (Rlang), Oktoberfest (Python), FragPipe(Java), Skyline (C#), or any other programming language, simplifying access from these languages for all currently implemented machine-learning models. Koina supports models from all major ML development frameworks. Currently, implemented models include Prosit, MS2Pip, DeepLC and AlphaPeptDeep. (b) A common peptide sequence interface was implemented for all models available via Koina to standardize pre- and post-processing steps based on a common input format, namely the PSI ProForma peptide notation standard, simplifying model comparisons.

We envision Koina to be an open-source, community-driven project. We welcome ML developers contributing to it by adding newly developed models to Koina, boosting their accessibility. The initial release of Koina features models focusing largely on the field of mass spectrometry-based proteomics, predicting a) gas phase fragmentation, b) chromatographic retention, and c) collisional-cross-section of peptides; specifically, Koina supports Prosit^12,17–19^, MS2PIP/DeepLC^11,20^, and AlphaPeptDeep^6^ (PeptDeep) (Fig 1a). The individual models were chosen because they are (1) among the most used ML/DL models used by the community (2) previously established by independent research groups, and (3) cover various modeling approaches, training data choices, and thereof, resulting in application limits. Together, they form a representative cross-section of the current peptide-property prediction landscape in proteomics. Providing an overview of state-of-the-art models available is one of the core functions of any model repository. Koina provides clear and concise documentation at https://koina.wilhelmlab.org/docs, covering all available ML models and significantly simplifying the discovery of novel machine-learning models. Notably, this documentation is semi-automatically generated based on an annotation file using OpenAPI and DOME standards. This means that ML developers adding new models to Koina do not need to be familiar with web development to provide easily accessible documentation for their models.

The Koina service encapsulates technically heterogeneous collections of models and makes them available through a single entry point that uses a common simple-to-use query interface (Fig 1b). This solves one of the main difficulties users encounter when using an ML model – a lack of documentation regarding the pre- and post-processing of input and outputs. Even when code is properly documented, most ML/DL developers use custom input output formats when developing their models; this unnecessarily complicates the work of scientists interested in using model predictions. Koina minimizes this hurdle for users and developers of ML/DL models by encapsulating pre- and post-processing together with the core model in a “workflow” or “execution graph” (Fig 2a). This simplifies the situation for users by abstracting a level of detail that is unnecessary for them. Since the execution graph supports Python code, ML developers can reuse the code they used during model development, allowing for parallelizing separate steps of the execution graph, vastly improving performance under heavy load. We have chosen the ProForma (Proteoform and Peptidoform Notation) notation format developed by the Proteomics standards initiative (PSI)^21^, meaning that tools and users that want to interface with Koina do not need to implement complicated file standards. This improves interoperability across various tools, allowing quick implementation of a Koina interface in various third-party applications.

**Figure 2:**
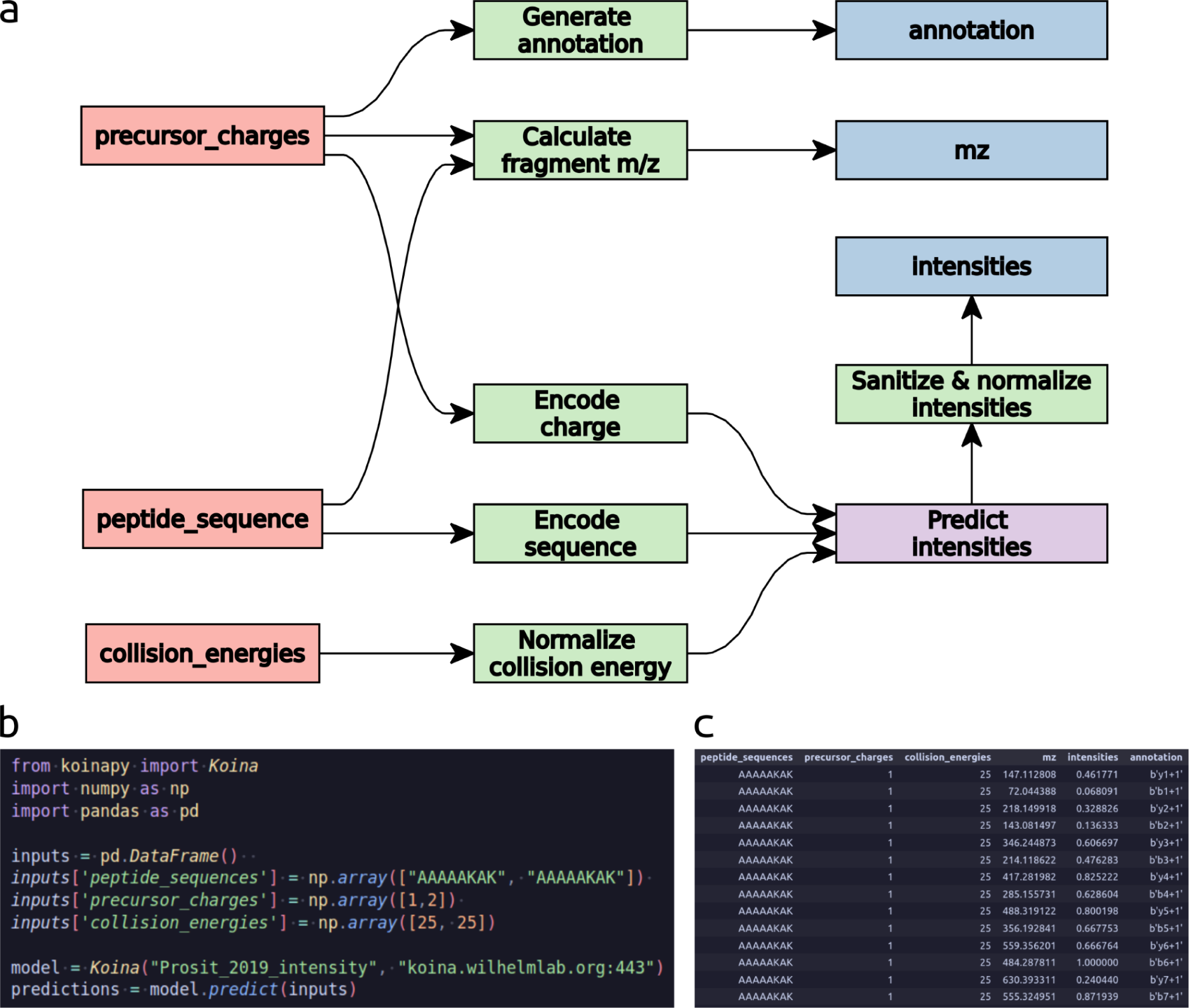
a) Execution graph for the Prosit_2019_intensity model. Inputs are colored yellow, (Python) pre- and post-processing scripts are colored green and turquoise, respectively. The TensorFlow neural network is purple. Outputs are blue. b) Code example for calling the Prosit_2019_intensity model using the KoinaPy client library as seen on koina.wilhelmlab.org. Simplifying interactions with Koina. c) Example output as generated by a call using KoinaPy.

Koina’s focus on encapsulating models and making them directly executable also ensures that all dependencies are explicitly encoded, ensuring long-term reusability. This is supported by a continuous integration (CI) pipeline using GitHub Actions. No changes to pre- and post-processing scripts, such as optimizing performance or updating dependencies, can have unintended effects on prediction reproducibility. Prediction reproducibility is supported by GitHub, where changes are transparently tracked, and separate docker images are released for every version. Version control is also supported through Zenodo, which is used to store binary model files and doesn’t allow changing files without creating a new version (Fig 1a). Different versions of the model can also be created. This allows most users to automatically use bug fixes while older model versions can still be accessed to ensure compatibility.

Koina abstracts most of the model related logic from the client by taking care of it on the server, the client only needs to handle some minimal processing in relation to sending HTTP requests, such as input formatting, batching, request preparation and error handling. To improve the availability of Koina further, we also developed client packages to trivialize connecting to Koina for the most common data science languages, Python (https://github.com/wilhelm-lab/koina) and R (https://github.com/wilhelm-lab/koinar). With these client packages, predictions can be generated in as little as four lines of code (Fig 2b/c). They can also serve as reference implementations for the development of client packages in other languages. For this, as well as other commonly used languages like Java, C#, and JavaScript, example code snippets are provided at koina.wilhelmlab.org that showcase how to fetch predictions for any model.

### Benchmarking prediction models for improved peptide identification with MSBooster and FragPipe

#### Koina is integrated into MSBooster and FragPipe

To increase Koina’s accessibility, we have integrated it into the FragPipe computational proteomics platform (https://fragpipe.nesvilab.org/). Specifically, the deep-learning-based MSBooster^4^ can query the server to gain access to many more models for prediction, allowing users to compare and mix-and-match models all within the same software environment. MSBooster formats batches of peptides and metadata into JSON files, and then sends these files to Koina via HTTP/S requests (Fig 3a). Koina then sends its predictions back in JSON format, and the results are parsed into MSBooster’s spectral library class to use later for PSM rescoring. Different prediction models can be specified, allowing users to change between DIA-NN^7^ (default in MSBooster in FragPipe) and the models supported on Koina. Models can be specified both via FragPipe (Supplemental Fig 1a) and standalone MSBooster (https://github.com/Nesvilab/MSBooster). Each model supports different kinds of PTMs and peptides (Table 1). Different models also run at different speeds (Fig 3b, Supplemental Data 1, Methods). DIA-NN runs the fastest, as it can be run locally, circumventing the time taken to transfer data to the Koina server and back. All the Koina models have comparable prediction times, slower than DIA-NN but still on the order of thousands of peptides per second. A plot showing the rate of Koina prediction is automatically generated (Supplemental Fig 1b).

**Figure 3.**
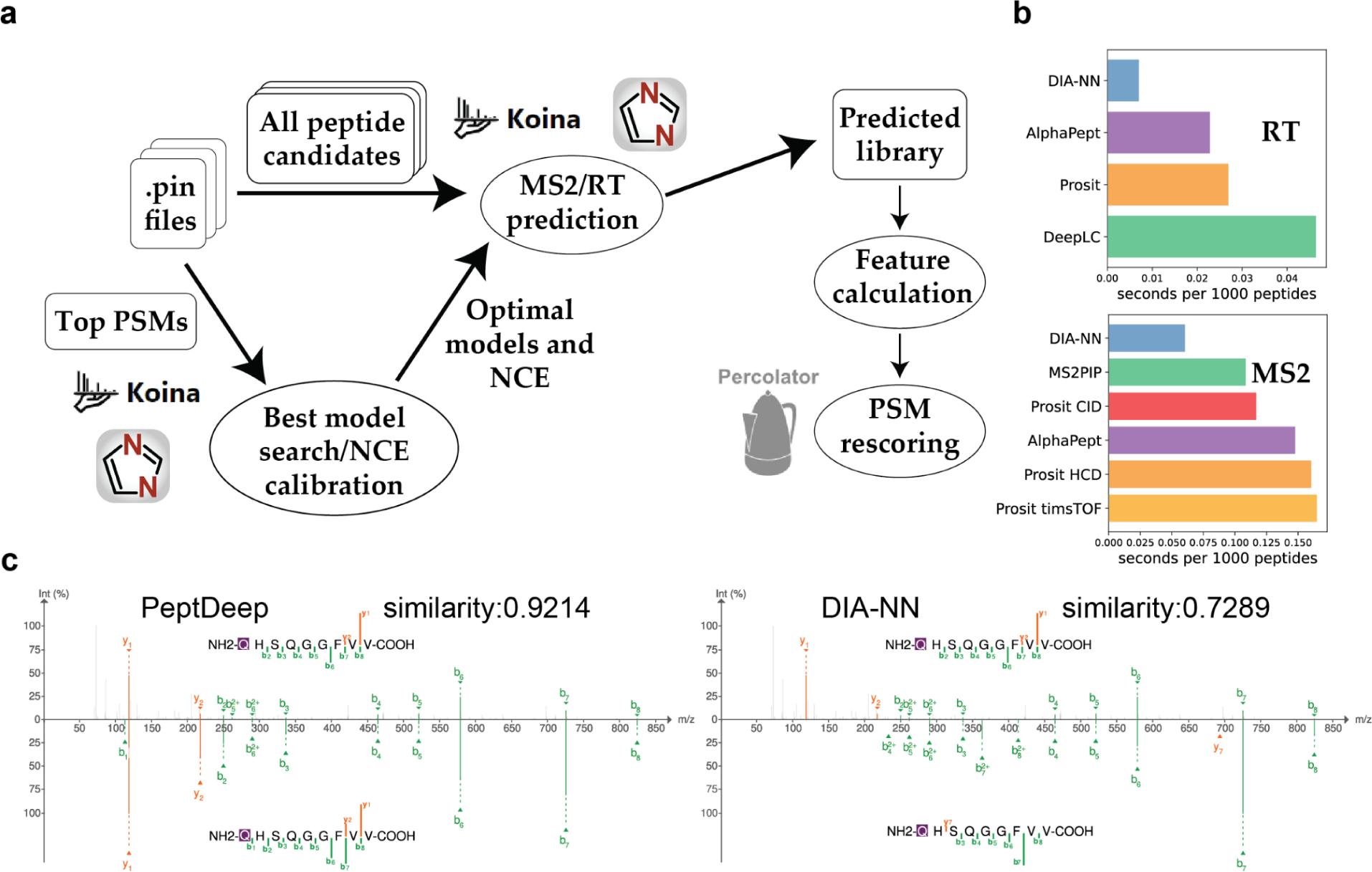
Koina integration in MSBooster in FragPipe. a) All peptide candidates are extracted from .pin files and predicted by either DIA-NN (available as part of FragPipe) or models available on Koina. These models can be specified manually, and NCE calibration is performed automatically if the model accepts NCE as a parameter. MSBooster can also use a heuristic algorithm to attempt to automatically choose the best performing MS2/RT model combination. The 1000 top PSMs are selected by default for the best model search and NCE calibration steps. b) Timing per 1000 peptides predicted was determined for models used in at least 2 experimental analyses. Each dataset was predicted ten times. c) Spectra 20200317_QE_HFX2_LC3_DDA_CM647_R01.42711.42711.2^26^ was visualized using either PeptDeep or DIA-NN predicted spectra, with the experimental spectra on the top and predicted spectra on the bottom. Unweighted spectral entropy was used as the MS2 similarity metric.

**Table 1.** Different models have different requirements for prediction. We consider if peptides longer than 30 amino acids (AA length > 30), phosphopeptides (Phospho), and peptides without N-terminal TMT labeling (n-term TMT optional) are allowed. A model was considered to support phosphopeptides not only if it could predict them with no error, but also if RT/fragment intensities changed when the PTM was specified.

|  | DIA-NN | Prosit | AlphaPeptDeep | MS2PIP | DeepLC |
| --- | --- | --- | --- | --- | --- |
| AA length > 30 | Yes | No | Yes | No | Yes |
| Phospho | Yes | No | Yes | No | Yes |
| n-term TMT optional | Yes | No |  |  |  |

Predicted libraries are saved as Mascot Generic Format (MGF) files if a Koina model was used, or as binary files if DIA-NN was used. These files can be reloaded if rerunning MSBooster from the command line to avoid calling the prediction model again. Either the MGF or binary file can be loaded into FragPipe-PDV^22^ for comparison of experimental and predicted spectra (Fig 3c). Here it becomes easier to visualize the differences between models’ predictions. For example, DIA-NN does not predict intensities for fragments shorter than three amino acids long, as they are less informative for DIA-NN’s peptide-centric approach. Fig 3c shows one example in which the pyroglutamated peptide Q[-17]HSQGGFVV exhibits a strong y1 fragment intensity which is matched by PeptDeep’s prediction but not by DIA-NN’s, resulting in a nearly 0.2 drop in spectral similarity, which ranges from 0 to 1.

One important piece of metadata required by some models is the normalized collision energy (NCE), which affects fragmentation patterns. NCE is an important parameter to tune based on the type of peptides being fragmented^23,24^. Indeed, other computational workflows have also noted the importance of calibrating NCE to maximize the similarity between experimental spectra and model predictions^1,12^. To account for differences between the learned NCE patterns of the predictors and NCE settings on individual instruments, MSBooster performs an NCE calibration step when using Koina models (Fig 3a). The top 1000 PSMs ranked by expectation value (evalue) across all pin files are extracted and predicted at all integer values in a certain range. Predicted and experimental spectra are compared with the unweighted spectral entropy similarity metric^4,25^, and the NCE value that produces the highest median similarity is selected when calling the model to predict all other peptide candidates from the pin files. A quality control plot showing the distributions of the similarity scores for the PSMs at each NCE value provides greater insight into this calibration step (Supplemental Fig 1c)

#### Phosphoproteomics data analysis

Having Koina connected to MSBooster enables users to compare model performance systematically and mix-and-match models. Because of differences in model architectures and training data, we hypothesize that the optimal model for rescoring will depend on the specifics of the dataset. To investigate this, we considered various types of data to pinpoint patterns between specific proteomics data types and optimal models. We first considered phosphoproteomics data. Along with DIA-NN, the only currently available Koina models that support phosphopeptide prediction are PeptDeep and DeepLC. We benchmarked model performance for two datasets. First, we processed phosphoproteome data from 30 different Arabidopsis thaliana tissues^27^ and counted how many phosphopeptides were identified. We found that PeptDeep outperformed DIA-NN both in terms of MSBooster’s spectral similarity and RT difference scores (Fig 4a). DIA-NN’s MS2 feature achieved an average improvement of 3.9% over baseline performance without MSBooster, while PeptDeep’s MS2 feature had a 4.1% improvement. PeptDeep’s superior MS/MS predictions are evident in Fig 4b, where its predictions for high-scoring “confident target” PSMs are concentrated closer to the maximum similarity of 1. Confident target PSMs are defined as those with evalues lower than the lowest evalue for a decoy PSM from the same pin file. The RT feature tells a similar story (DIA-NN 1.8% improvement over baseline, DeepLC 2.3%, PeptDeep 3.7%) (Fig 4a). It is less apparent from the distributions of the RT difference feature which RT model performs best (Fig 4e, Supplemental Fig 2). DIA-NN’s RT predictions improved identifications the least in the Arabidopsis dataset, yet its high-scoring target PSMs have the smallest RT differences. However, DIA-NN also assigned decoy PSMs smaller RT differences compared to the other models, which decreased target-decoy separation and ultimately led Percolator to find the RT feature less informative. Because PeptDeep’s MS2 and RT predictions surpassed DIA-NN’s for phosphopeptides, we hypothesized that the two features in combination, when calculated using PeptDeep’s predictions, would also perform better than when using DIA-NN’s predictions. We confirmed this, finding a 4.6% increase over baseline using DIA-NN and 6.0% increase using PeptDeep (Fig 4a). This is in line with the magnitude of improvement expected for a closed MSFragger search on tryptic DDA data^4^.

**Figure 4.**
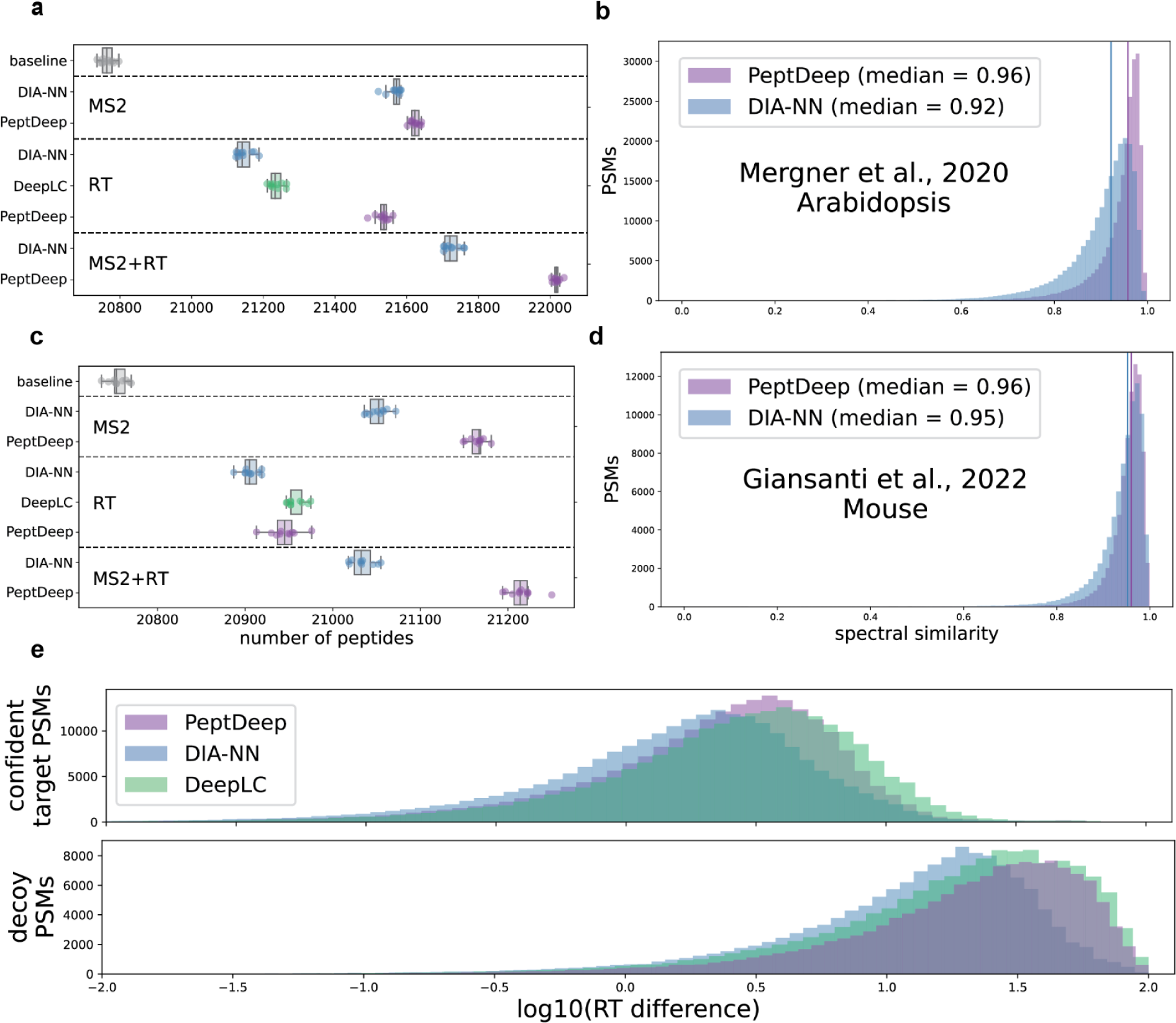
Phosphoproteomics rescoring with various models. (a) Numbers of peptides identified using different models on the Arabidopsis thaliana data across 30 tissues^27^. Each model was repeated ten times, each with a different Percolator random seed. “Baseline” is the peptides identified when excluding MSBooster before Percolator rescoring. MSBooster calculated and added the unweighted spectral entropy feature (MS2), delta RT loess feature (RT), or both (MS2+RT). (b) The distribution of spectral similarity scores for confident target PSMs (those PSMs with lower expectation values or “evalues” than the lowest evalue assigned to a decoy PSM in the same pin file). The features were calculated using predictions either from PeptDeep or DIA-NN. (c-d) The same as (a-b) but for the 8 mouse pancreatic ductal adenocarcinoma cell lines^28^. (e) The distribution of delta RT loess scores for confident target and decoy phosphorylated PSMs from the Arabidopsis thaliana dataset. The log10 of each value plus a small pseudocount was added for clearer visualization of the distribution differences. Features were calculated using predictions from DIA-NN, PeptDeep, or DeepLC.

Comparison of the three models on 8 mouse pancreatic ductal adenocarcinoma (mPDAC) cell lines^28^ showed similar results (Fig 4c-d). Interestingly, DeepLC performed similarly to PeptDeep here, identifying 20958 and 20945 phosphopeptides on average, respectively. Based on these findings, we believe that PeptDeep should be used instead of DIA-NN for phosphoproteomics analysis.

#### Prosit performs the best across multiple HLA experiments

We and others have shown that human leukocyte antigen (HLA) immunopeptidomics data benefits greatly from PSM rescoring with predicted libraries^3,6,17,18,29–31^, largely in part because the increased nonspecific peptide search space leads to many more spurious matches to decoy sequences. HLA peptides fall into one of two classes based on which class of major histocompatibility complex (MHC) molecule they bind, similarly designated class I or II. They have been widely studied because of increased interest in HLA peptides for use as biomarkers and in immunotherapeutics ^32,33^. Recent developments in data-independent acquisition (DIA) and instrumentation (such as the development of the Bruker timsTOF machine) have provided deeper and more reproducible views of the HLA peptidome ^34,35^. We benchmarked all models on a variety of HLA datasets ^26,36–38^ (Fig 5a-g, Supplemental Data 2). Overall, the DIA-NN and Prosit RT models performed the best over baseline, while the Prosit models dominated in the MS/MS category. In addition, we found that combining the best RT and MS/MS models outperformed the current FragPipe default of using DIA-NN’s predictions to compute both the MS2 and RT features for all datasets except for one of the DIA datasets (Fig 5e). This improved ability to identify peptides provides a compelling reason for incorporating the many models on Koina into FragPipe, as models trained on large nontryptic datasets can give better results than DIA-NN when creating predicted spectral libraries.

**Figure 5.**
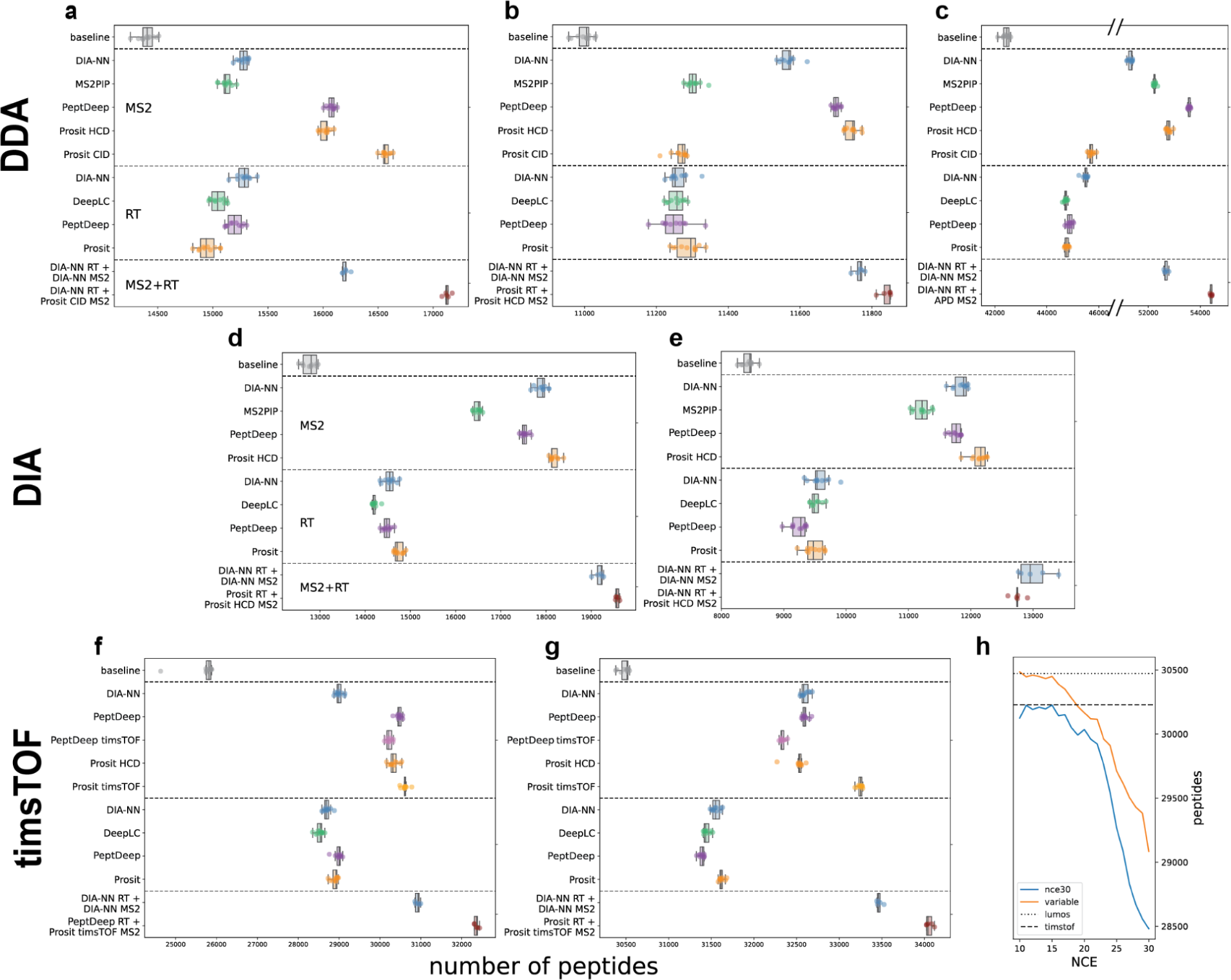
HLA rescoring with various models. The datasets used are (a-b) Marcu et al., 2021 class I and II^36^, (c-d) Pak et al., 2021 class I DDA and DIA^26^, (e) Ritz et al., 2017 class I DIA^37^, and (f-g) Phulphagar et al., 2023 class I and II on a timsTOF single-cell proteomics system^38^. “PeptDeep timsTOF” is the PeptDeep model with “timsTOF” set as its instrument metadata, while “PeptDeep” used either “Lumos” or “QE” as its instrument metadata, depending on what was listed in the mzML files or manuscript text of the respective datasets. Hatch marks in the x-axis indicate sections of the plot removed due to empty space, unoccupied by any peptide counts. (h) The number of peptides identified by the nce30 and variable PeptDeep transfer learned models at various eV values in the class I data represented in (f). Each eV value was tested three times using three different Percolator random seeds. The dashed lines for “lumos” and “timsTOF” indicate the average peptides identified by the PeptDeep and PeptDeep timsTOF models respectively in (f).

The findings here highlight how specialized models outperform generic ones. First, we note the differences in performance when rescoring spectra from different fragmentation methods. Distinct higher-energy collision dissociation (HCD) and collision induced dissociation (CID) models exist for Prosit and MS2PIP. However, only the HCD MS2PIP model is currently available on Koina, while the CID model is on the MS2PIP server at https://iomics.ugent.be/ms2pip/^11^. Therefore, our comparison is limited to using the Prosit models. Marcu et al.^36^ used CID to fragment precursors in their class I sample (Fig 5a). We found that the Prosit CID model performed best here, achieving 15% more peptide identifications over baseline and also outperforming its Prosit HCD counterpart. Likewise, the Prosit HCD model performed better on datasets using HCD fragmentation^26,36^ (Fig 5b-c). Whichever model is superior is also evident in the distribution of target PSMs’ spectral similarity score distribution, which is shifted to lower similarity when the wrong fragmentation model is used (Supplemental Fig 3).

Specialized models improve rescoring MS/MS spectra generated not only from different fragmentation methods, but also from different mass spectrometers. Strangely, though Prosit timsTOF improves over the Prosit HCD model trained on Orbitrap data for both class I and II data from Phulphagar et al.^38^, this does not extend to the PeptDeep models, where “Lumos” PeptDeep predictions helped identify more peptides than “timsTOF” PeptDeep predictions did (Fig 5f-g). We believe the reason for this discrepancy is in how PeptDeep and Prosit were trained on spectra of different NCEs. While Prosit timsTOF was trained on energies ranging from 20-70 eV, PeptDeep timsTOF was only trained on 32-52 eV, according to the training data listed in their supplemental data^6^. Furthermore, all timsTOF spectra were annotated as 30 eV during the PeptDeep training phase. Though spectra do differ across Orbitrap and timsTOF instruments, there are certain hotspots across energy levels where they are highly correlated^18,39^. Specifically, precursors fragmented with lower energies (around 20% and 20 eV on Orbitrap and timsTOF instruments, respectively) produce nearly identical spectra across instruments. Phulphagar et al. employed a scheme where precursor ion mobility and fragmentation energy were inversely correlated, with energy ranging from 55-10 eV. Though MSBooster performs an NCE calibration step to find the best NCE value for spectral prediction, the lower bound of this experimental NCE range is far below that of what PeptDeep timsTOF was trained on, potentially resulting in suboptimal predictions. However, because low energy fragmented spectra are similar between Orbitrap and timsTOF, the PeptDeep model predicting for a Lumos instrument performs better, since it was trained on spectra with HCD energy as low as 20%. Indeed, when we rescored another dataset acquired on a timsTOF with similar ion mobility settings as the data PeptDeep was trained on^40^, both PeptDeep and Prosit timsTOF models outperformed their Orbitrap counterparts (Supplemental Fig 4).

To confirm our hypothesis that annotating spectra in the training set with the correct NCE would improve PeptDeep timsTOF performance, we transfer-learned new PeptDeep models using the same datasets Prosit timsTOF was trained on and tested their performance across a range of low collision energy values on the class I timsTOF data from Phulphagar et al.^38^ (Fig 5h, Methods). Models were trained in one of two ways. The first, named “nce30”, was trained by annotating all PSMs as 30eV, as was implemented in PeptDeep’s timsTOF training. By excluding informative NCE metadata in the training, this transfer-learned model was still unable to achieve the number of peptides identified by the PeptDeep Lumos model, despite having observed the fragmentation of many new non-tryptic peptides. The second model was dubbed the “variable” model and used the NCE information provided for each peptide. Here, we were able to identify similar peptide numbers to the Lumos model as early as 15eV, yet the variable model was still unable to surpass the Lumos model’s performance. We conclude that encoding all training spectra as 30eV resulted in PeptDeep timsTOF learning an average representation of spectra across collision energies that did not extrapolate well to lower energies; when it is properly encoded, NCE influences peptide spectral predictions strongly enough to result in noticeable changes in the final numbers of peptides identified, though we were unable to train a PeptDeep timsTOF model that outperformed what was learned by the Lumos model.

Interestingly, we also note that class I datasets benefited more from predictions than class II datasets did, in line with previous literature^17^ (Fig 5a-c, f-g, Supplemental Data 2). This is evident across all RT and MS/MS models. A potential reason for this decreased improvement is that longer peptides are more difficult to predict, likely because of the increased number of peptide bond fragmentations possible.

#### Astral DIA data analysis

Recently, the Orbitrap Astral mass spectrometer (Thermo Scientific) has piqued the interest of the proteomics community for its wide dynamic range, sensitivity of detecting low abundance precursors, and accurate and precise quantification^41,42^. The higher acquisition rates compared to Orbitrap analyzers allow for narrower isolation windows, producing spectra of reduced complexity in data independent acquisition (DIA) without sacrificing throughput. With the rising interest in narrow window DIA (nDIA), we analyzed two short LC-gradient datasets using 2 Th isolation windows using an MSFragger-DIA based workflow in FragPipe (see Methods). The first dataset included three technical replicate injections of Hap1 human cell lysate^43^. As there are currently no Astral-specific models on Koina, we investigated whether Orbitrap or timsTOF models showed better performance. We found that for both Prosit and PeptDeep, the Orbitrap mode performed better on this dataset, though only by a small margin (Prosit: 60280 vs 60118; PeptDeep: 60586 vs 59716) (Supplemental Fig 5a). Regardless, DIA-NN, which is instrument-agnostic, performed the best both for MS2 and RT features. DeepLC identified fewer peptides than average compared to DIA-NN’s RT module, though the difference was insignificant (57645 vs 57673 peptides; two-sided t-test, p>0.05). DIA-NN’s MS2 and RT features also performed the best when combined. They achieved a 27.3% increase in peptide identifications over baseline. Meanwhile, combining the next best features of DeepLC for RT and PeptDeep for MS/MS only achieved a 25.5% increase over baseline. Similar findings were obtained from the second dataset of 46 fractionations in technical triplicates of HEK293 cells^44^ (Supplemental Fig 5b). Here, using DIA-NN for both features increased peptides identified by 18.5% and outperformed PeptDeep MS2 and DeepLC RT (17.9%). This lower percent increase in identifications may be attributed to the fractionated dataset’s already deeper proteome coverage at baseline, identifying more than 3 times the number of peptides as the first dataset. From these analyses, we find that Astral nDIA experiments may benefit most simply from DIA-NN predictions, though this leaves room for improvement if an Astral-specific model is trained.

#### DIA-NN and Prosit TMT models perform comparably

Currently, the only two models available in FragPipe for TMT-based prediction are DIA-NN and Prosit, both of which were trained on hundreds of thousands of sequences^19^. We examined two datasets to see whether these models differed in not only the number of peptides identified, but also in their quantification quality. We first analyzed 36 fractions of one plex of TMT11-labeled peptides from a study of lung adenocarcinoma (LUAD) tumors^45^. On this dataset, Prosit outperformed DIA-NN’s MS/MS predictions, but not by a significant amount (93306 vs 93269 TMT-labeled peptides, t-test: p>0.05) (Fig 6a). Meanwhile, DIA-NN greatly outperformed Prosit’s RT predictions (93877 vs 92787, t-test: p<0.05). Using both DIA-NN features improves over baseline by 3.7%, in contrast to the 2.8% improvement with Prosit. Using DIA-NN for RT and Prosit for MS2 prediction, we identified 94991 peptides on average, a 3.8% improvement over baseline and statistically significantly more than using DIA-NN to predict both peptide properties (t-test: p<0.01). Combining the DIA-NN and Prosit models is enabled by MSBooster’s access to DIA-NN and Koina.

**Figure 6.**
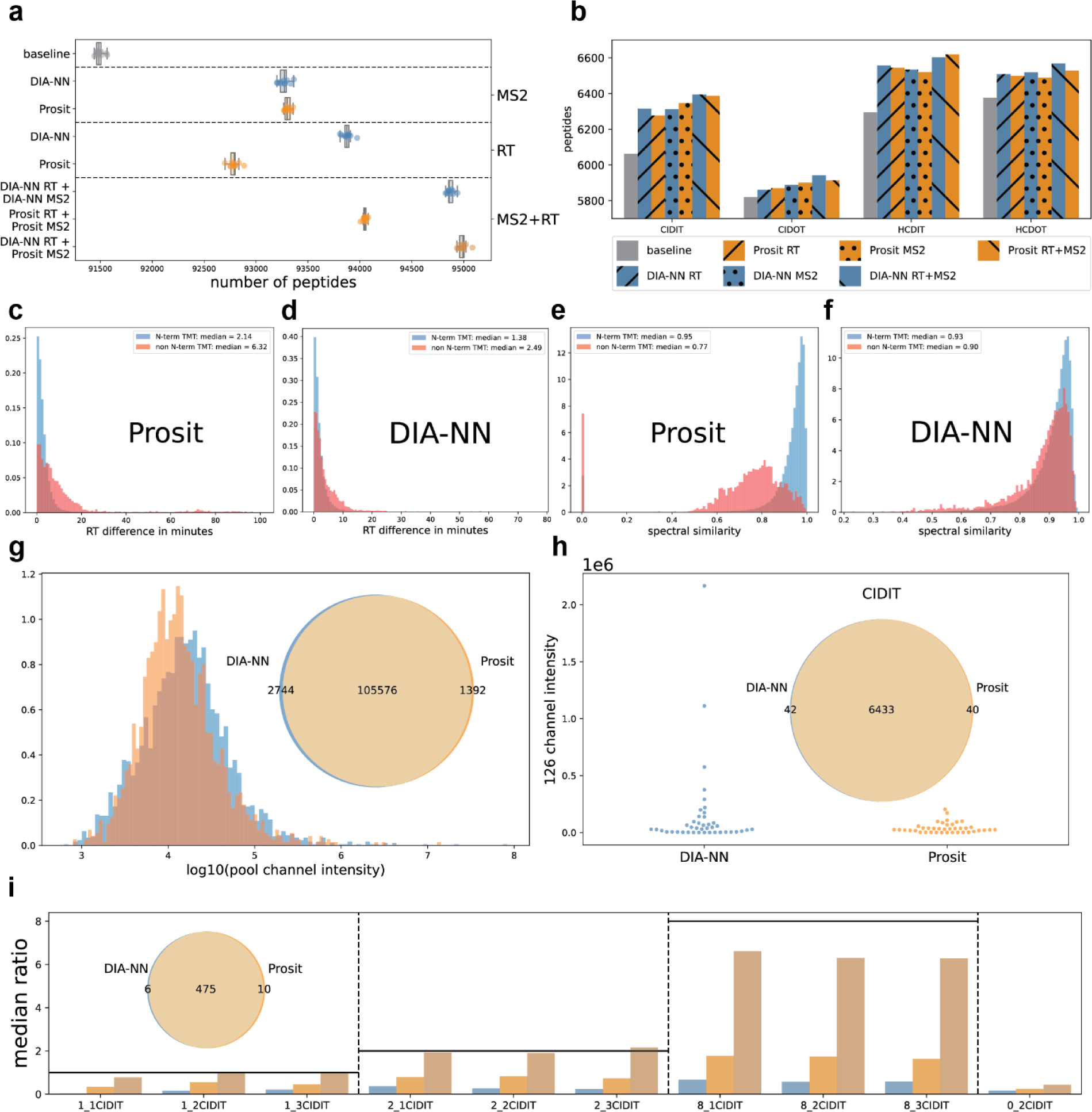
Comparison of TMT models. (a) Number of TMT11-labeled peptides identified with 10 iterations of PSM rescoring. Samples were 36 fractions of the LUAD dataset^45^. (b) TMT11 peptides identified in a yeast/human proteome mixture^19^. 4 combinations of settings were produced by matching CID/HCD fraction with orbitrap (OT) or ion trap (IT) analyzers. Runs using DIA-NN’s predictions are in blue and runs using Prosit’s are in orange, with separate hatch marks designating whether RT, MS2, or both feature types were used. (c-f) Histograms depicting feature scores for confident target PSMs separated by N-terminally labeled (blue) or not (red). c-d depicts the RT difference score, while e-f depicts the spectral similarity score. (g) 2744 and 1392 unique TMT PSMs were identified by DIA-NN and Prosit. The 126 channel was a pooled sample and non-zero reporter ion intensities from this channel are visualized here. (h) Unique PSMs from each model in the CID IT run were selected and the reporter ion intensities in the 126 channel were visualized in a swamplot. Both human and yeast PSMs were included here. (i) Quantified yeast peptides from DIA-NN and Prosit were compared via Venn diagram. For each subset of this Venn diagram, the median ratio of the peptides in each channel to the reference 126 TMT channel (not shown here) is visualized, with a horizontal black line indicating what the expected ratio is.

DIA-NN is more flexible than Prosit in that it does not assume peptides are TMT-labeled at the N-terminus (Table 1). To adapt Prosit predictions to accommodate peptides lacking N-terminal TMT, we simply assign them the RTs of the same peptide sequences but with N-terminal TMT and shift the m/z of their b-ions by the TMT label’s mass while retaining the same intensities. Neither model considers over-labeling of TMT on serine, so MSBooster uses the strategy above to create predictions for peptides with labeled serine for both models. To explore how well this strategy produces predictions for peptides lacking N-terminal TMT, we compared MSBooster feature scores between peptides without and without the N-terminal label in a subset of high-ranking confident target PSMs, 2.3% of which were unlabeled at the N-terminus. We found that while both DIA-NN and Prosit exhibit higher accuracy for N-terminally labeled peptides, Prosit exhibited more noticeable differences between the two groups both in terms of the RT difference and MS2 similarity features (Fig 6c-f, Supplemental Fig 6a-b). In addition, Prosit does not support prediction for peptides longer than length 30. MSBooster assigns spectral similarity values of 0 and predicted indexed RTs (iRTs) of 0 to PSMs without peptide predictions, further penalizing Prosit’s performance; this is most clearly seen in Fig 6e with a large amount of confident target PSMs having spectral similarities of 0, and in Supplemental Fig 6a with the cloud of PSMs straying far from the RT calibration curve. In contrast to this “regular” search, we ran a “restricted” database search, setting N-terminal TMT as a fixed modification and limiting the peptide digest length to 30, to determine if the two models performed more similarly without these biases. Without MSBooster’s features, the baseline number of TMT-labeled peptides decreased by more than 2000 (Supplemental Fig 6c). Again, we found that DIA-NN performed best for the RT feature and Prosit for the MS2 feature, though the gap in RT feature performance decreased (Prosit regular vs restricted: 1.4% vs 2.0% increase over baseline; DIA-NN: 2.6% vs 2.7%) and the gap in MS2 feature performance increased (Prosit regular vs restricted: 2.0% vs 2.0%; DIA-NN: 1.9% vs 1.7%). This suggests that when N-terminal TMT labeling is incomplete or when many longer peptides are expected, the flexibility of DIA-NN is beneficial.

When analyzing a separate TMT11-labeled human-yeast protein mixture^19^, our initial findings that DIA-NN was best for RT and Prosit for MS/MS did not hold. This dataset had four settings combining either HCD or CID fragmentation with Orbitrap (OT) or ion trap (IT) analyzers. Here we found that DIA-NN and Prosit performed comparably in the number of TMT peptides identified (Fig 6b). Interestingly, though Prosit TMT takes the fragmentation mode (HCD or CID) into account during prediction and DIA-NN does not (nor was it trained on any CID data), the models performed comparably on CID data. N-terminal labeling rates for confident target PSMs were 0.05%, 0.04%, 0.04%, and 0.05% for CID IT, CID OT, HCD IT, and HCD OT, respectively.

We next assessed whether the unique PSMs identified by the two models differed by their reporter ion intensities. In the LUAD dataset, DIA-NN found 2744 unique TMT PSMs to Prosit’s 1392 (Fig 6g). We compared the pooled channel’s MS2-based reporter ion intensities between the unique sets and found a statistically significant difference between their median intensities (DIA-NN: 1.58e4, Prosit: 1.22e4; p<0.01, Mann-Whitney U test). Given the wide dynamic range of reporter ion intensities from all the PSMs in this dataset, this difference does not seem particularly meaningful. Indeed, none of the four MS3-based acquisition methods in the yeast dataset showed a statistically significant difference between DIA-NN and Prosit’s quantification (Fig 6h, Supplemental Fig 7) (p>0.01, Mann-Whitney U test).

Finally, we wished to check the validity of the unique peptides from each model. The yeast dataset provides a good benchmark because each channel has a known amount of yeast protein spiked into a constant amount of human protein, resulting in the expected ratios of 1:1:1:2:2:2:8:8:8:0 for yeast proteins in channels 127N to 131C compared to the 126 channel. Therefore, we focused on peptides originating from the yeast proteome to assess quantification accuracy. In the data acquired by CID IT, the two models had 475 quantified yeast peptides in common (Fig 6i). DIA-NN and Prosit had 6 and 10 unique quantified peptides, respectively. The shared, DIA-NN-specific, and Prosit-specific peptide sets all showed increasing median ratios across the triplicate channels, with the shared set most closely following that expected 1:2:8 ratio. The average median ratio for DIA-NN was 0.13, 0.29, and 0.61; for Prosit it was 0.45, 0.78, and 1.71; for the shared set it was 0.93, 1.91, and 6.40. Similar trends exist in the three other acquisition methods (Supplemental Fig 8-9). Though peptides solely identified by Prosit seemed to exhibit higher median ratios than those from DIA-NN, these differences were not statistically significant (p>0.01, Mann-Whitney U test). The unique peptides displayed much lower median ratios than what was expected. While some of these peptides may be false positives, most have high spectral similarities, suggesting that these are real identifications (Supplemental Fig 10). Potential explanations for these reduced ratios include TMT reporter intensity suppression and insufficient unique peptides in each group making it difficult to achieve an accurate median ratio. Overall, most yeast peptides found by DIA-NN and Prosit were overlapping, and many of the peptides that are unique to each model are likely valid. We do not find sufficient evidence that these unique sets are different in any meaningful way. Therefore, we suggest that these models can be used interchangeably, but DIA-NN should be favored when there are many PSMs matched to peptides that are longer than 30 amino acids or with unlabeled N-termini.

Automated heuristic model search accurately determines the best model combination In the above sections, we presented empirically which models work best for phosphorylation, HLA, Astral DIA, and TMT data. However, it is outside the scope of this manuscript to exhaustively consider all types of proteomics data, and even within the same type of data there is variability in the best-performing models, as can be seen in the HLA datasets considered here (Fig 5a-g, Supplemental Data 2). Furthermore, as Koina continues to grow and support more models, it may become overwhelming for users to determine which combinations of models best improves their PSM rescoring. To maximize numbers of identified peptides, we have incorporated an optional module in MSBooster that attempts to quickly determine the best MS/MS and RT model for a dataset. Importantly, the selected models may be from different frameworks (e.g. DeepLC for RT and Prosit for MS/MS), a benefit conferred by Koina’s one-stop shop for all these models.

Rather than having to rescore the entire dataset using each available model, we implemented a heuristic approach that only gets predictions for the top 1000 PSMs (with no consideration of their status as targets or decoys) and selects models based on the agreement between their predicted and experimental values. When developing this algorithm, we initially chose the MS2 model with the highest median similarity and RT model with the lowest median RT difference, but we found that the algorithm would sometimes report models we did not find empirically performed the best in our tests. Figure 7 depicts one such case for an HLA DDA class I dataset ^36^. Our algorithm chose Prosit rather than DIA-NN as the best RT model (Fig 7b). Though Prosit had the lowest median delta RT of 1.38 minutes and DIA-NN the largest of 2.03 minutes out of the models tested, it was actually DIA-NN’s RT predictions that identified the most peptides on average, with 6.1% increase over baseline. Compare this to Prosit RT which performed the worst with its 3.8% improvement over baseline and 312 fewer peptides than DIA-NN. This is a similar finding to Fig 4e, where DIA-NN assigned confident PSMs the lowest delta RT scores out of the RT prediction models, yet still identified the fewest peptides. Supplemental Fig 11 shows this same analysis for the other datasets.

**Figure 7.**
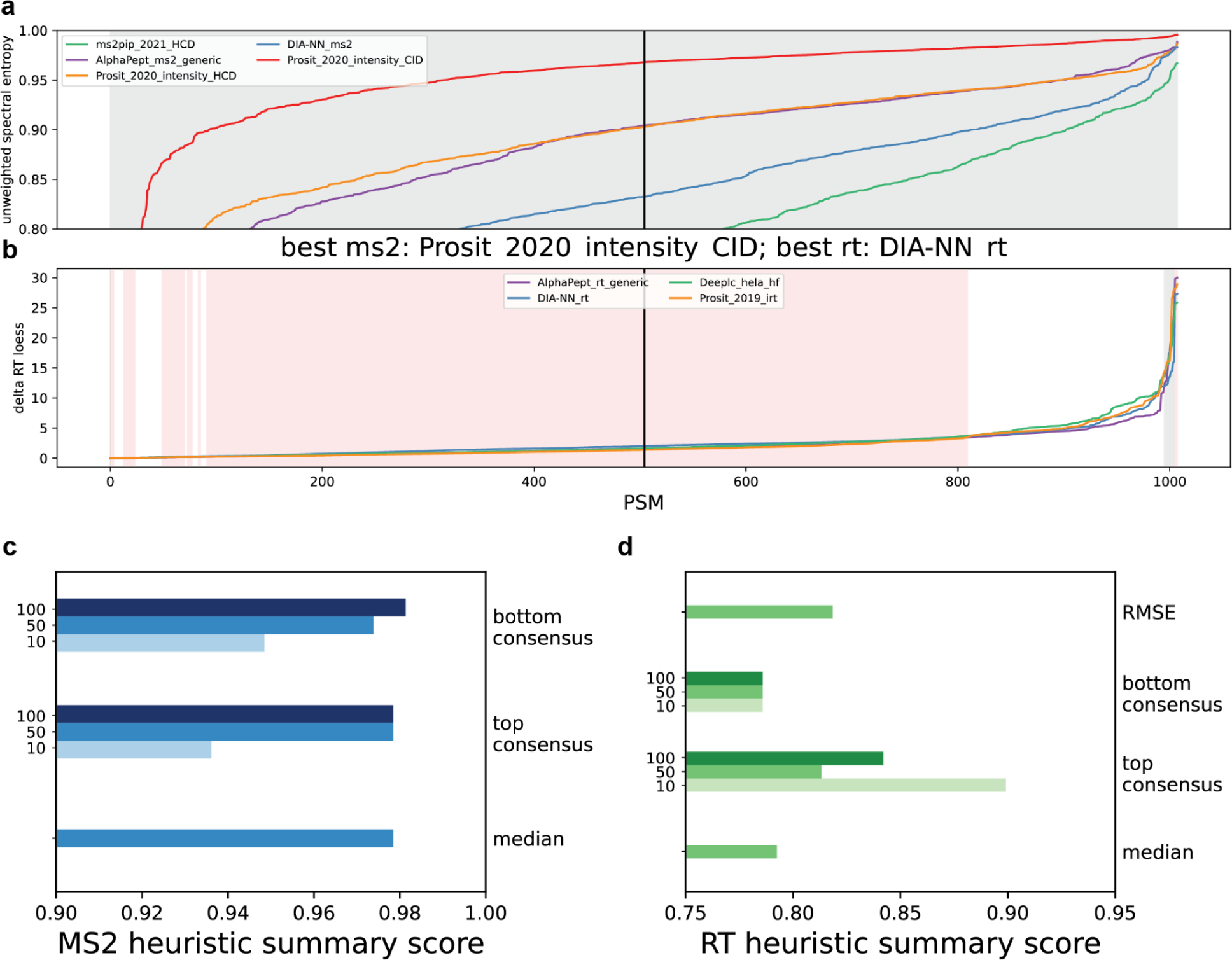
Heuristic best model search. Predicted MS/MS spectra (a) and RT values (b) for the top 1000 PSMs of an HLA DDA class I dataset^36^ were compared to their observed values. The calculated spectral similarity and RT difference scores were ordered in increasing order and plotted for each model. The black line indicates the position of the median value. Gray regions indicate positions at which the model that empirically performed the best (Fig 5a) would be selected by the heuristic. Red regions indicate positions at which the model selected by the heuristic does especially worse than the empirically best model (i.e. one standard deviation above the mean performance of the heuristic-selected model is less than one standard deviation below the mean performance of the empirically best model). The performance of multiple heuristic methods was summarized across all datasets for MS2 models (c) and RT models (d). For the top and bottom consensus methods, the values of N PSMs tested were 10, 50, and 100 PSMs. Greater detail explaining how the summary score was calculated is provided in the Methods section.

It was clear that the algorithm could be improved, so we tested several other metrics and summarized their performance across all datasets used here (Fig 7c-d, Methods). We found our “top consensus” metric performed well for finding the optimal RT model. The top consensus metric sorts the 1000 PSMs in increasing feature value (higher MS2 similarity or greater RT difference) and takes the N highest feature values reported by each model. At each index in these sorted sublists of length N, the algorithm casts a vote for which model is preferable (highest MS2 similarity and smallest RT difference at that specific index). Whichever model has the highest number of votes is chosen. We found that taking the 10 largest delta RT values was better able to determine the best-performing RT model compared to the median method. Specifically, the top 10 consensus method achieved an overall score of 0.899, meaning the heuristic search suggested models across our testing datasets that on average identified 89.9% of the peptide numbers compared to a method that is able to find the empirically best RT model all the time; compare this to the median method, which achieved a score of 0.793. While the best MS2 method was taking the bottom 100 lowest spectral similarity scores (score of 0.981, Methods), the median similarity method performed very similarly (score of 0.978), and the methods chose different models in only two out of the thirteen dataset. Therefore, there was insufficient justification to move away from the median method, so we continue to use it for determining the best MS2 model.

We have now implemented this improved model determination metric in the algorithm, but users can adjust it to their needs by making use of command-line MSBooster’s ability to exclude models from the best model search which they deem unnecessary (e.g. timsTOF models should not be used for Orbitrap data and vice versa, and only models supporting phosphorylation prediction should be used in phosphoproteomics analyses).

### Conclusion & Outlook

This study demonstrates how Koina simplifies access to machine learning models in the proteomics domain by providing a unified platform. Multiple popular ML models, covering different frameworks and approaches, have been implemented to predict a) gas phase fragmentation, b) chromatographic retention, and c) collisional-cross-section of peptides. We encourage ML developers to submit their models to Koina, which will not only enhance the platform’s diversity but also broaden the impact and accessibility of their models.

The common model interface implemented for Koina enables third-party tool developers to integrate access to any model available through Koina. This can serve as a starting point for developing a new standard to define common input and output formats of ML models in the proteomics domain, further improving interoperability. An important issue for discussion is the handling of modifications absent in the training data. Currently, this varies between models. Prosit’s strategy limits predictions to evaluated modifications, enhancing accuracy but reducing applicability. Conversely, MS2PIP disregards modifications not in the training data and provides the best estimate of the fragment ion intensity as if the amino acid were unmodified, adjusting only fragment peak m/z.

The currently available public Koina instances (koina.wilhelmlab.org) make it easily accessible to new users. The distributed design of Koina allows scaling to community needs by enabling heavy users to dedicate hardware to support it. Alternatively, the provided docker image facilitates the easy deployment and scaling of private instances, which is helpful for cases involving confidential data or performance-critical tasks.

We illustrate how Koina’s integration in FragPipe and MSBooster enables benchmarking of a variety of state-of-the-art models without having to run each model in a separate analysis pipeline. We gained insights into which models perform well on specific data types and touched on important considerations for training and evaluating these models. Notably, we found that PeptDeep sometimes performed suboptimally on timsTOF data, while Prosit did not encounter this issue. This aligns with findings of a recent study that evaluated strategies for training peptide-property models with scarce data^46^, suggesting that multi-task learning performs worse than single task learning. PeptDeep provides a single model covering fragment intensity prediction of multiple instrument types. Prosit, on the other hand, crafts separate models for different instruments. This allows for their models to learn different weights that can better accommodate different instruments, unlike the PeptDeep approach in which the majority of parameters are shared between instruments.

Additionally, we developed a heuristic best model search module in MSBooster, anticipating the future availability of many additional models. Interestingly, we observed that calculating the median feature score for the top 1000 PSMs from a dataset does not always result in identifying the empirically best-performing model. Even if finding the optimal model can not be guaranteed, the newly developed improved scoring method demonstrated high accuracy, picking RT and MS/MS models that are on average 90% and 98% as good as the best performing models, respectively. High-accuracy predictions remain crucial, but our findings indicate that they do not always translate to the most significant gains in peptide identifications. Optimizing predicted spectral libraries for PSM rescoring is a complicated question that may benefit from examining deep-learning-based feature scores for high-scoring targets and decoys, as well as those PSMs on the border of the false discovery rate cutoff. Future investigation into this can further improve our heuristic search algorithm, contributing to the continuous improvement of ML predictors in the proteomics domain.

In summary, Koina offers a robust platform for simplifying access to diverse ML models in proteomics, promoting interoperability and ease of integration. Our study highlights the strengths and limitations of existing models, providing a foundation for future enhancements. By inviting developers to contribute to Koina, we aim to foster a collaborative environment that propels advancements in machine learning for proteomics, ultimately benefiting the entire scientific community.

## Methods

### Koina Infrastructure

Koina models are hosted using Nvidia Triton, an inference server supporting ML models developed in all major prediction ML development frameworks. To unify the interface of Koina and to make ML models usable without knowledge of pre- and post-processing, Python models are used to transform inputs/outputs. Models are tied together with the ensemble model functionality, which is referred to as an execution graph within the context of this manuscript. Models are made available by the Kserve API implemented by Nvidia Triton Inference Server, providing access using both REST and gRPC interfaces. Nginx is used for load balancing of the public Koina network. A docker image is provided to simplify the deployment of new Koina instances. To minimize its size, models are not stored in the image itself but rather dynamically fetched from Zenodo once the server is deployed. Both the image as well as the OpenAPI documentation provided at koina.wilhelmlab.org are automatically created, tested and deployed using a custom GitHub Actions workflow.

### FragPipe analysis and database searches

All analyses were done using FragPipe 21.0 with MSFragger 4.0^47^ for database searching, Percolator 3.6.4^48^ for PSM rescoring, ProteinProphet^49^ for protein assignment, Philosopher 5.1.0^50^ for FDR filtering and reporting, and IonQuant 1.15.0^51^ and TMT-Integrator 5.0.7^52^ for TMT quantification and summarization. MSBooster 1.2.2^4^ was used to turn deep-learning predictions into features before PSM rescoring, while MSBooster 1.2.30 was used for timing and to demonstrate heuristic model searching. Databases for searches were downloaded from UniProt, and all included common contaminants and reversed sequence decoys. The human database was downloaded March 18, 2022; mouse December 19, 2023; Arabidopsis June 2016 (the fasta available from Mergner et al., 2020; and yeast April 15, 2024. A yeast-human database was created by combining the two individual databases and used to search data from Gabriel et al., 2022. All workflows were adapted from those available in FragPipe, with specific workflows and parameters for each tool provided in https://github.com/Nesvilab/MSBooster/tree/master/Koina%20manuscript%20resource.

### Model timing within MSBooster

To make timing of models robust, we calculated the slope of a linear regression correlating the number of peptides submitted for prediction to the total prediction time. We only timed models that were used in at least two datasets. Each dataset was predicted ten times. 256G of RAM and 55 threads were used for DIA-NN prediction and Koina asynchronous prediction.

### Heuristic best model search scoring

Multiple methods were developed to approximate the best model for a dataset without having to rescore all PSMs and process them through downstream tools. First, the 1000/P PSMs with the lowest expectation values are selected from each pin file processed by MSBooster in the run, where P is the number of pin files. Next, all PSMs are sorted in ascending order of their unweighted spectral entropy (MS/MS) or delta RT loess (RT) score. Then, each heuristic method was applied.

- Median: The median score calculated using the predictions of each model is determined. The MS2 model with highest median similarity and RT model with lowest RT difference are chosen as the best models.
- Top consensus: N PSMs with the largest values are selected. Largest values for the PSM features when testing MS/MS models mean those with the highest spectral similarity; when testing RT models, large values mean the greatest deviations from the RT calibration curve. The values of N tested are 10, 50, and 100. At every position from 1 to N, the model with the best PSM feature value (greatest MS2 similarity or smallest RT deviation) cast a vote. The model with the most votes was selected.
- Bottom consensus: This method is similar to the top consensus method, but it focuses on the PSMs with lowest MS/MS similarity and smallest RT deviation.
- RMSE: The root mean squared error of RT deviations.

Once each heuristic method had produced its pick for best model, the average number of peptides identified empirically across 10 runs with different Percolator random seeds was divided by the average peptides identified by the model with the highest average. This was applied separately to RT and MS2 models. This ratio of peptides identified by the heuristically chosen model to peptides identified by the empirically best model was averaged across multiple datasets to produce the final score shown in the figures. For phosphoproteomics datasets, all peptides were considered, not only phosphopeptides. For TMT datasets, all peptides were considered, not just those quantified.

### Transfer learning PeptDeep and assessing model performance across an NCE range

Transfer learning a timsTOF model was accomplished with peptdeep 0.2.0. Nontryptic data from Adams et al. was downloaded from PXD043844 and their reanalysis of tryptic data was downloaded from MSV000092462. Each PSM (peptide sequence, fragment masses, and fragment intensities) was extracted from the hdf5 files from PXD043844. PeptDeep requires certain formatting of the input data for training, including MaxQuant msms.txt files. Only charge 1 fragments were listed in these msms.txt files. Fragments and intensities from the hdf5 files, which include charge 2 fragments, were used to replace those listed in the msms.txt files for each PSM, as designated by a distinct raw file/scan number pair.

Two models were trained. The “nce30” model was trained as described in the PeptDeep supplementary data^6^, where all PSMs were labeled as 30eV for timsTOF training. The “variable” model was trained using the average collision energies extracted from accumulatedMsmsScans.txt files. Both models were trained for 20 epochs with batch size of 1024, learning rate of 1e-4, 5 warm-up epochs and dropout of 0.1.

We predicted spectral libraries using the NCE30 and variable models from 10-30eV. Input files containing the peptides to predict were generated by MSBooster. The resulting libraries were saved in mgf files and provided to MSBooster to add the spectral similarity score to the pin files for HLA class I timsTOF data^38^. Downstream analysis by Percolator, ProteinProphet, and Philosopher filtering was repeated thrice for each NCE.

## Acknowledgments / Funding

We thank the for hosting a Koina instance FGCZ ETHZ|UZH and Marco Schmidt for configuring the networking. Furthermore, we would like to thank the organizers of the EuBIC-MS - Developers Meeting 2023 for giving us a chance to start this project there. We also thank all members of Wilhelm and Nesvizhskii Labs for their valuable input and feedback.

This research was in part funded by the Munich Data Science Institute (L.L.); European Union’s Horizon 2020 Program under Grant Agreement 823839 [H2020-INFRAIA-2018-1; EPIC-XS] (W.G.); an ERC Starting Grant [101077037] (E.K., M.W., W.G.); National Institutes of Health grants R01-GM-094231, U24-CA210967, and U24-CA271037 (K.L.Y., F.Y., K.L., A.I.N.); and the Proteogenomics of Cancer Training Program 5T32-CA140044-12 (K.L.Y., A.I.N).

## Data availability

Source code for Koina is available at https://github.com/wilhelm-lab/koina. Documentation for Koina is available at https://koina.wilhelmlab.org/. The source code for the R-client package is available at https://github.com/wilhelm-lab/koinar. Source code for MSBooster is available at https://github.com/Nesvilab/MSBooster. FragPipe is available for download at https://fragpipe.nesvilab.org/.

## Notes

### Competing Interest Statement

M.W. is a co-founder and shareholder of MSAID GmbH and OmicScouts GmbH, with no operational role in both companies. The remaining authors declare no competing interests.

