## Supplemental Figures for "Koina: Democratizing machine learning for proteomics research"

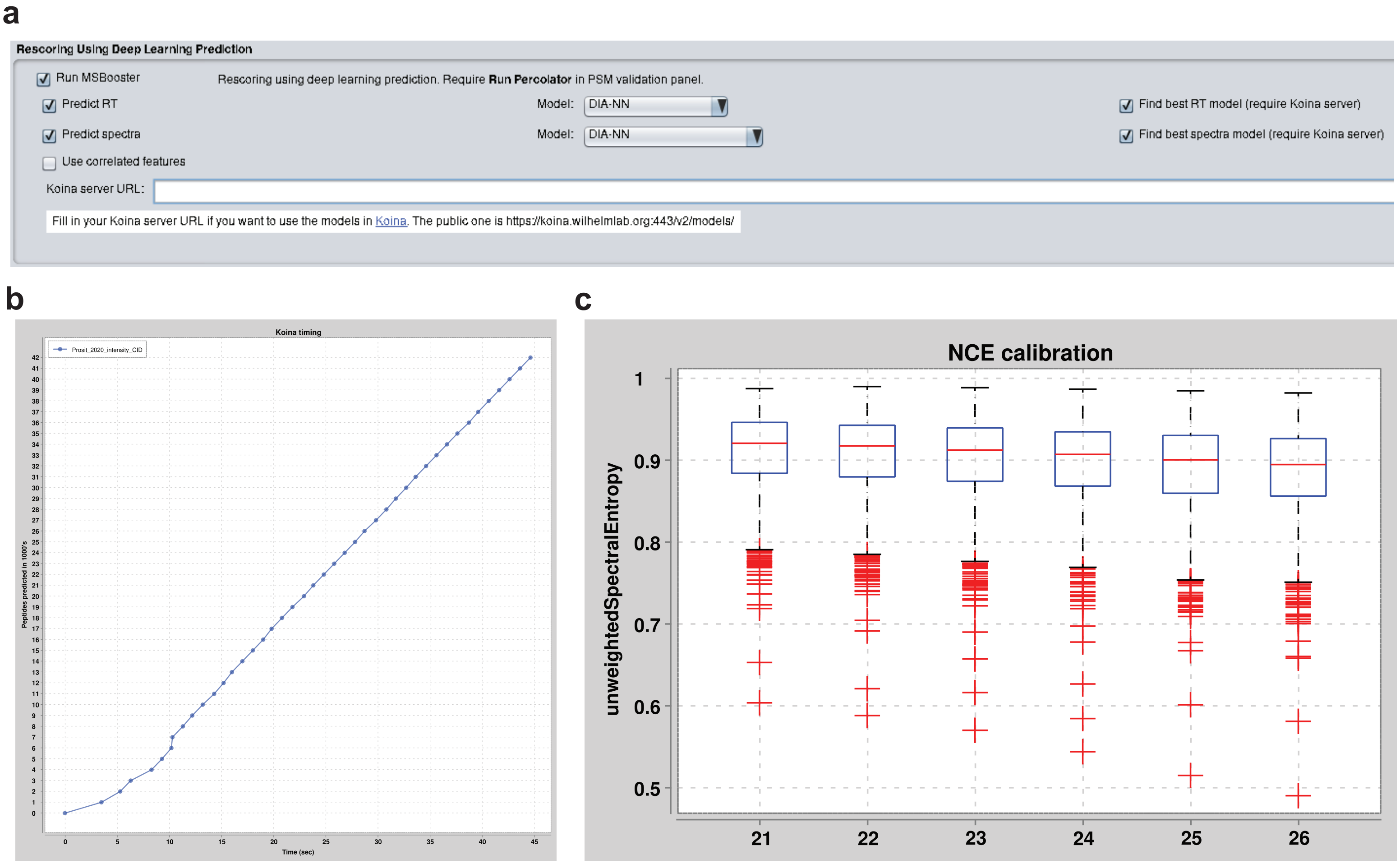


Supplemental Figure 1. (a) Screenshot of the MSBooster section under the FragPipe Validation tab. RT and spectral features can be used together or separately, with a dropdown menu to choose a model. If “Find best RT/spectra model” is checked, then a heuristic algorithm is enabled to attempt to determine the combination of models that maximizes peptides identified. One can also manually set an RT model but enable search for the best spectra model, and vice versa. A textbox is included to set the path to the Koina server, which may be a private, locally hosted one, or the public one provided. (b) Peptides are submitted in batches of 1000 to the Koina server. Total time taken is recorded on the x-axis. (c) Similarity box-and-whisker plot across sequential normalized collision energy (NCE) values for the PSMs collection during the NCE calibration step. The top whisker is the third quartile plus 1.5 times the interquartile range.

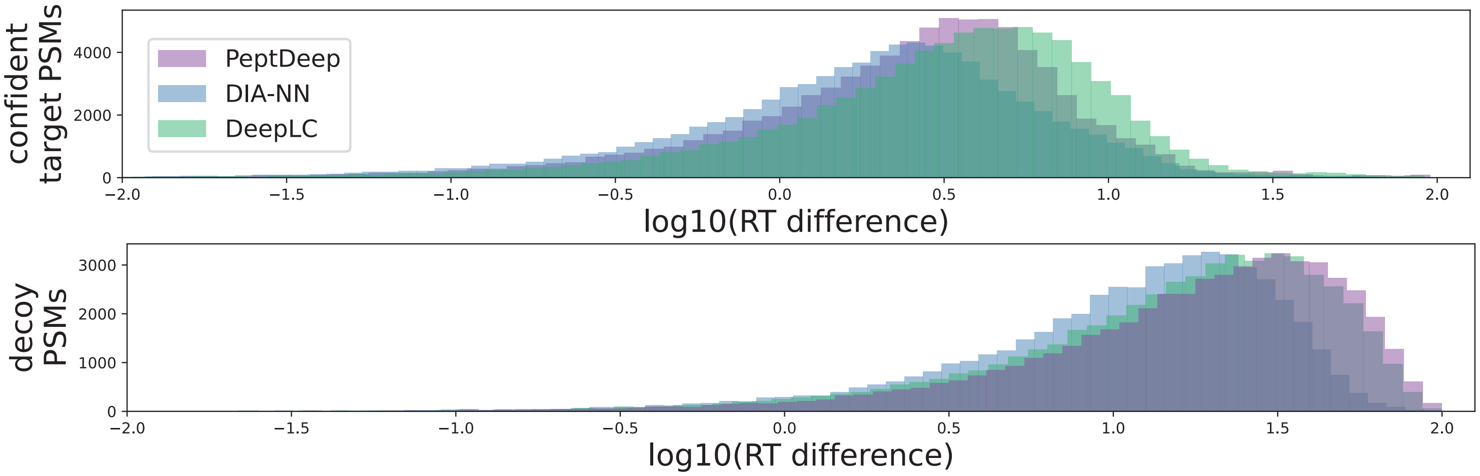


Supplemental Figure 2. The distribution of delta RT loess scores for confident target and decoy phosphorylated PSMs from the mouse PDAC dataset (Giansanti et al., 2022). The log10 of each value plus a small pseudocount was added for clearer visualization of the distribution differences. Features were calculated using predictions from DIA-NN, PeptDeep, or DeepLC.

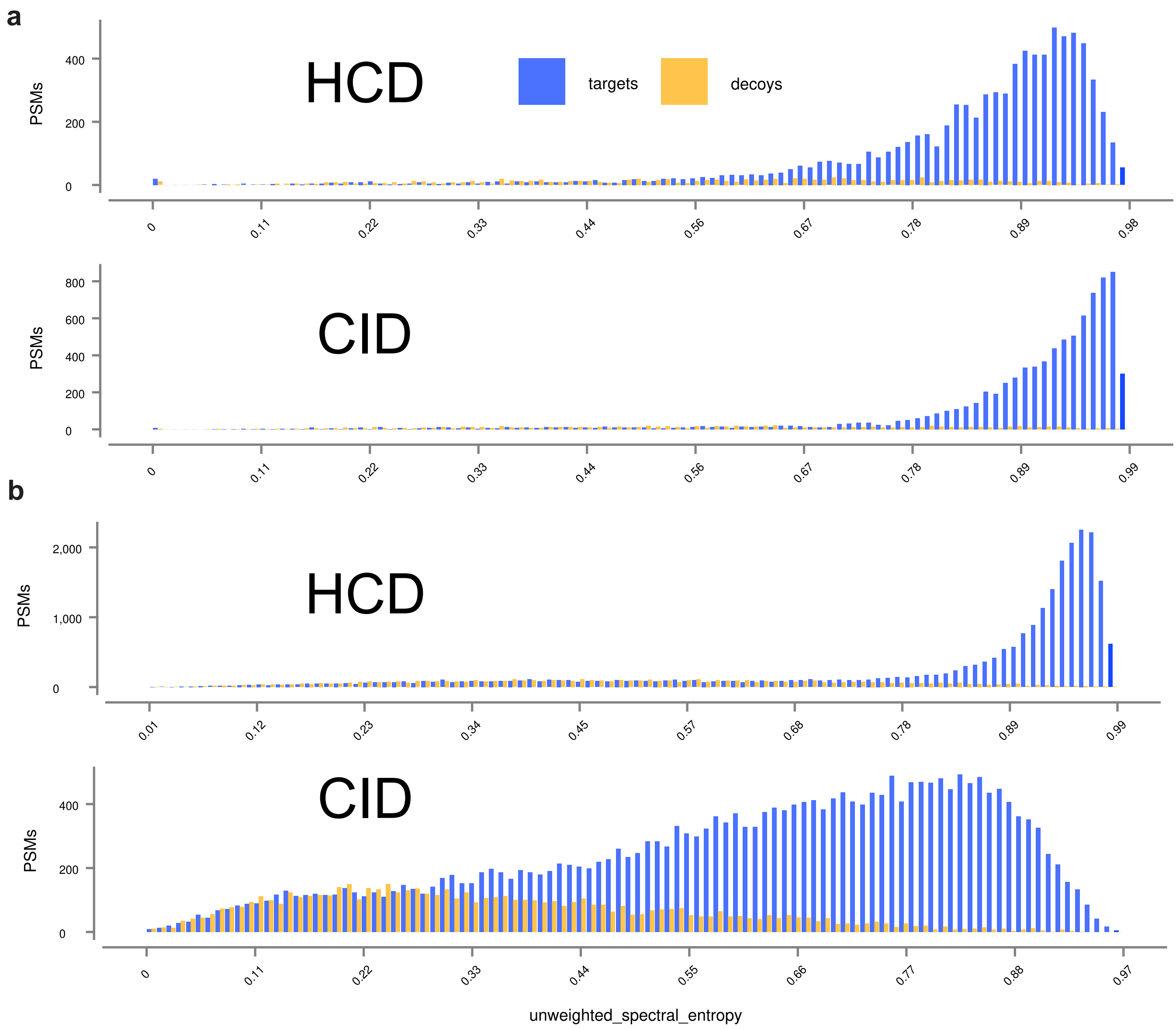


Supplemental Figure 3. Target and decoy PSM distributions for the unweighted spectral entropy MS/MS similarity feature when comparing experimental fragment ion intensities with predicted intensities from the Prosit_2020_intensity HCD and CID models. Distributions are shown for PSMs from one pin file each from a) Marcu et al., 2021 and b) Pak et al., 2021.

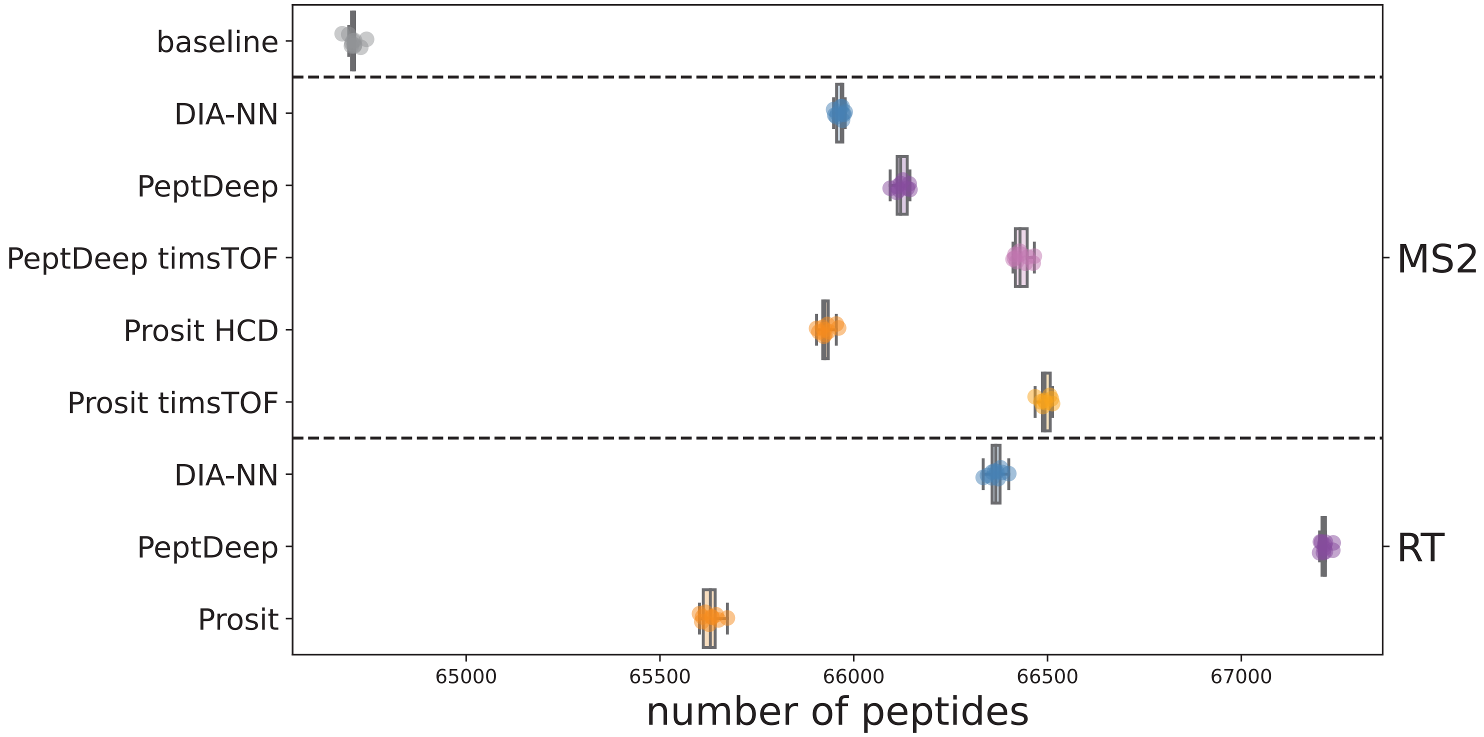


Supplemental Figure 4. Swarmplot and box-and-whisker plot of the peptides identified in data collected on a timsTOF mass spectrometer from Meier et al., 2018.

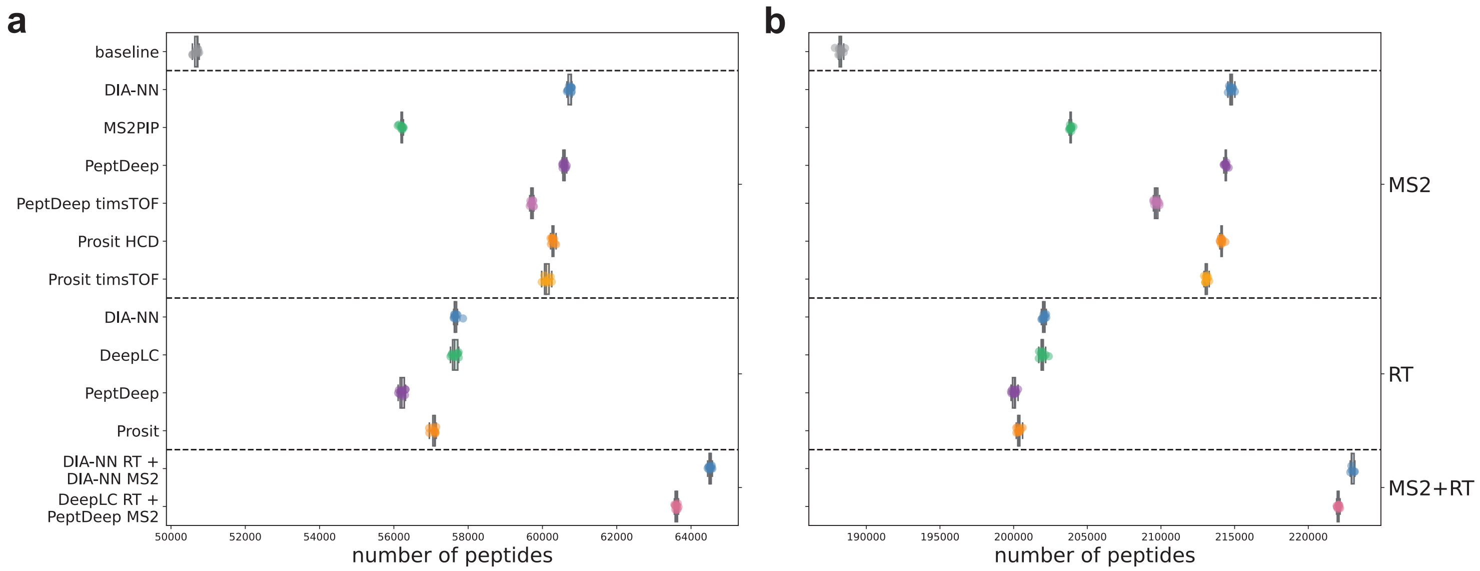


Supplemental Figure 5. Swarmplot and box-and-whisker plot of the peptides identified in data collected on Orbitrap Astral instruments from Serrano et al., 2024 (a) and Guzman et al., 2024 (b).

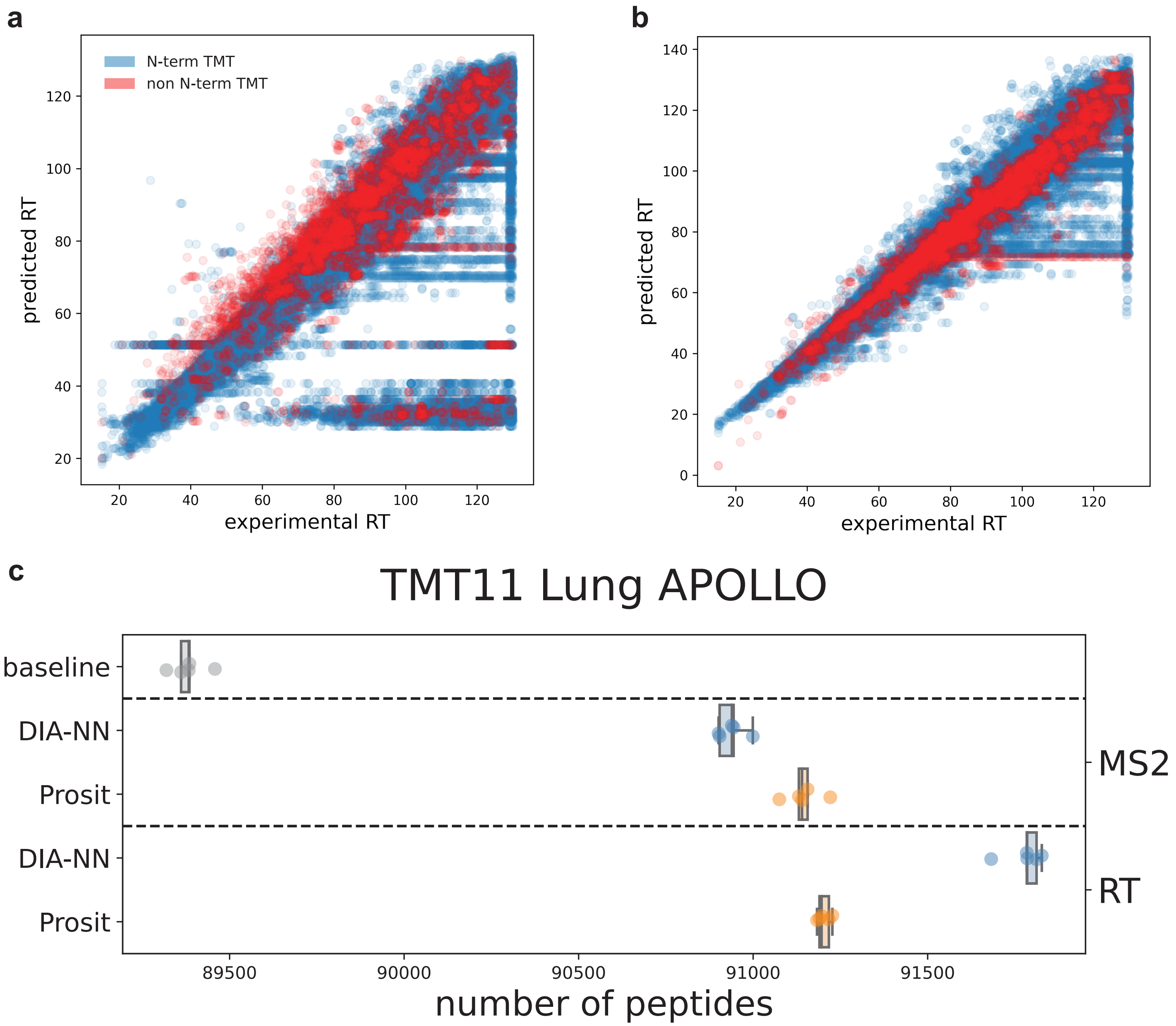


Supplemental Figure 6. (a-b) The correlation between experimental and predicted RTs in minutes. Peptides labeled with (blue) and without (red) TMT at the N-terminus are plotted for Prosit (a) and DIA-NN (b). (c) The number of TMT peptides identified when database search is "restricted". In-silico digest was done only up to 30 amino acids long, and N-terminal TMT was a fixed modification.

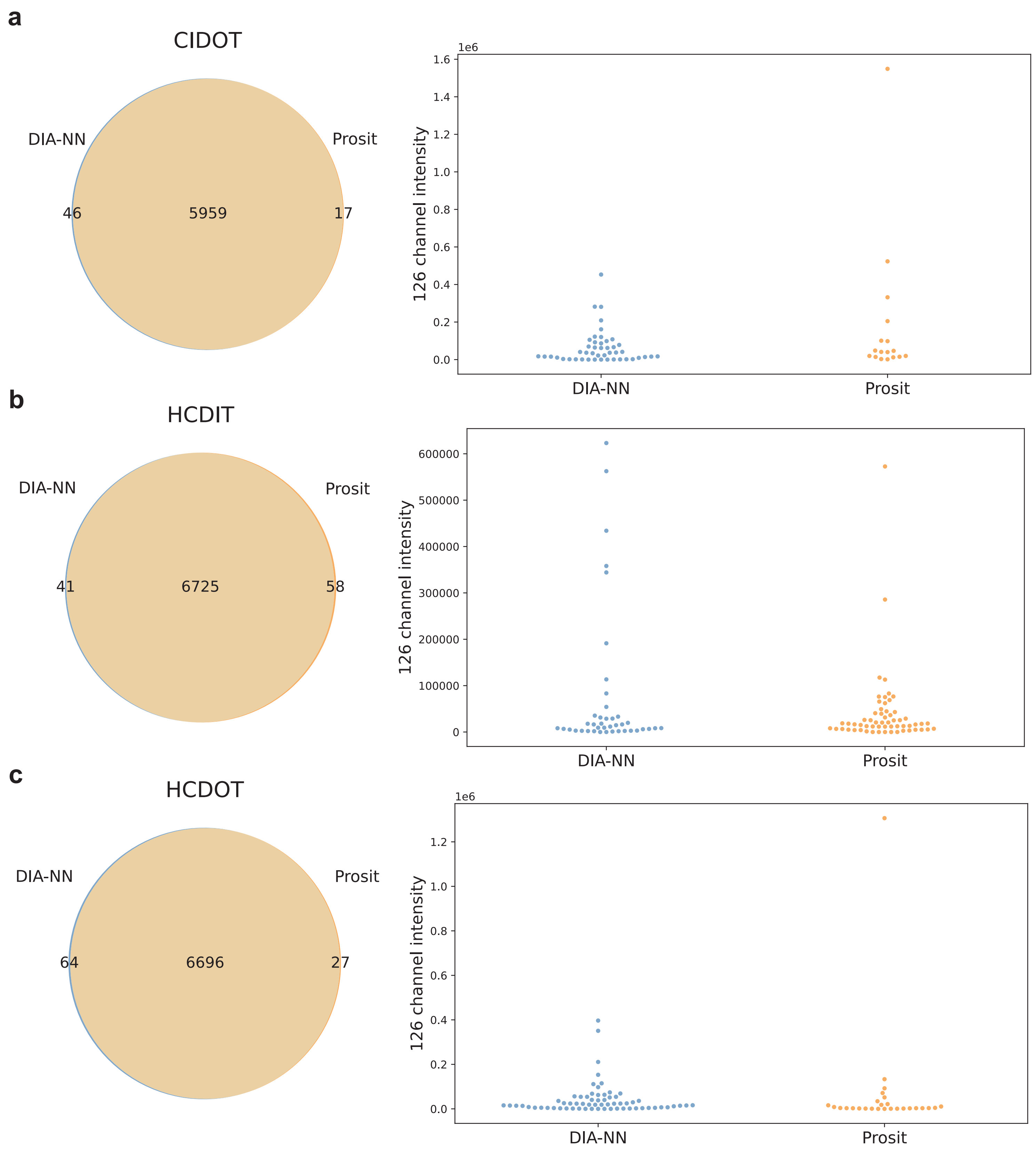


Supplemental Figure 7. Venn diagrams show the ovelap of PSMs that pass FDR filtering with DIA-NN or Prosit. The reporter ion intensities of the 126 channel (reference channel of HeLa-yeast 10:1 mixture) for these unique PSMs is also plotted as a swarmplot.

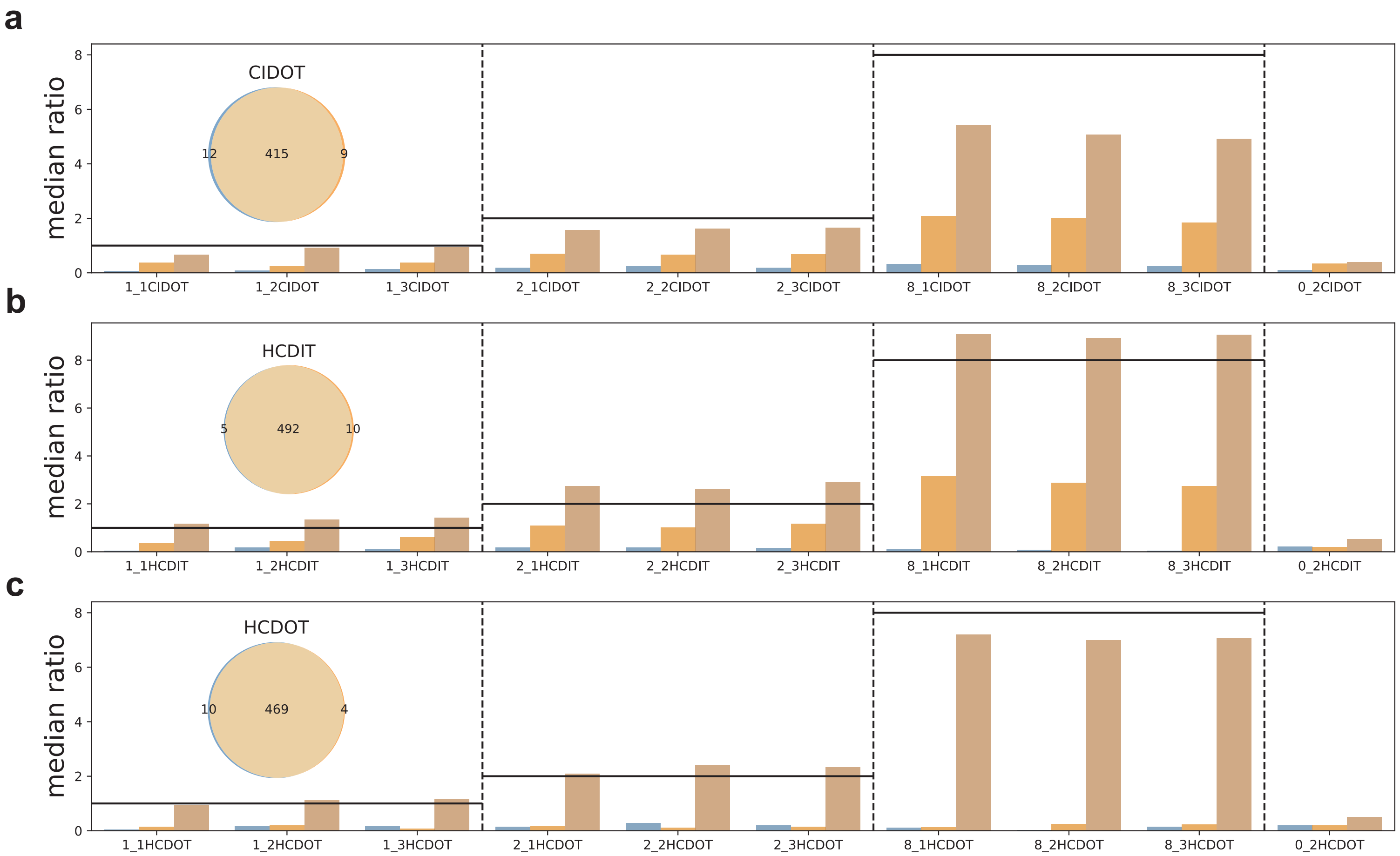


Supplemental Figure 8. Venn diagrams show the ovelap of yeast peptides from DIA-NN (blue) and Prosit (orange). The median ratios of each channel to the 126 channel for the subsets of the Venn diagram are plotted, with the black line showing the expected ratio.

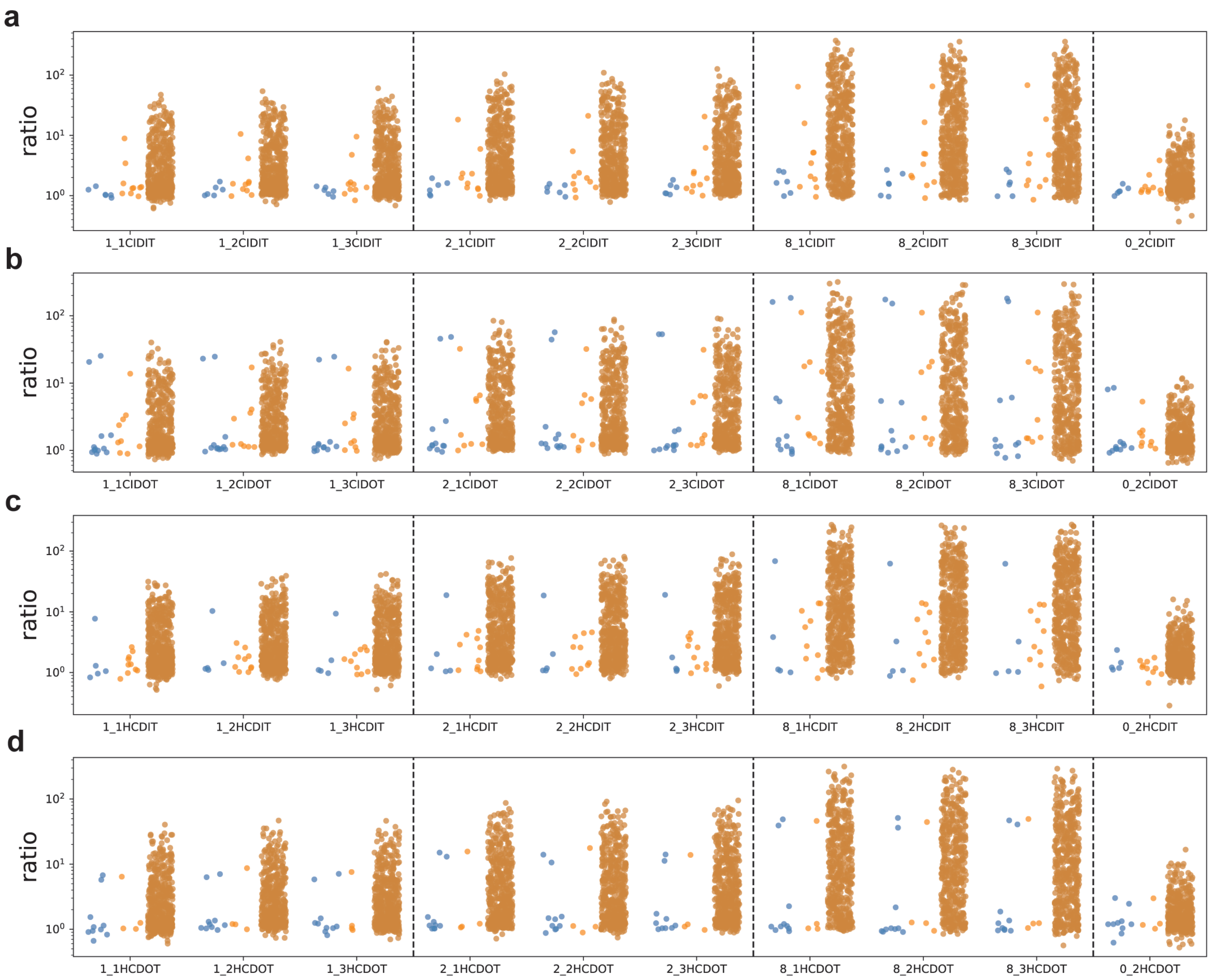


Supplemental Figure 9. The subsets from the Venn diagrams (Fig 4i, Supplemental Fig 8) are used here. Every dot represent a peptide quantification ratio.

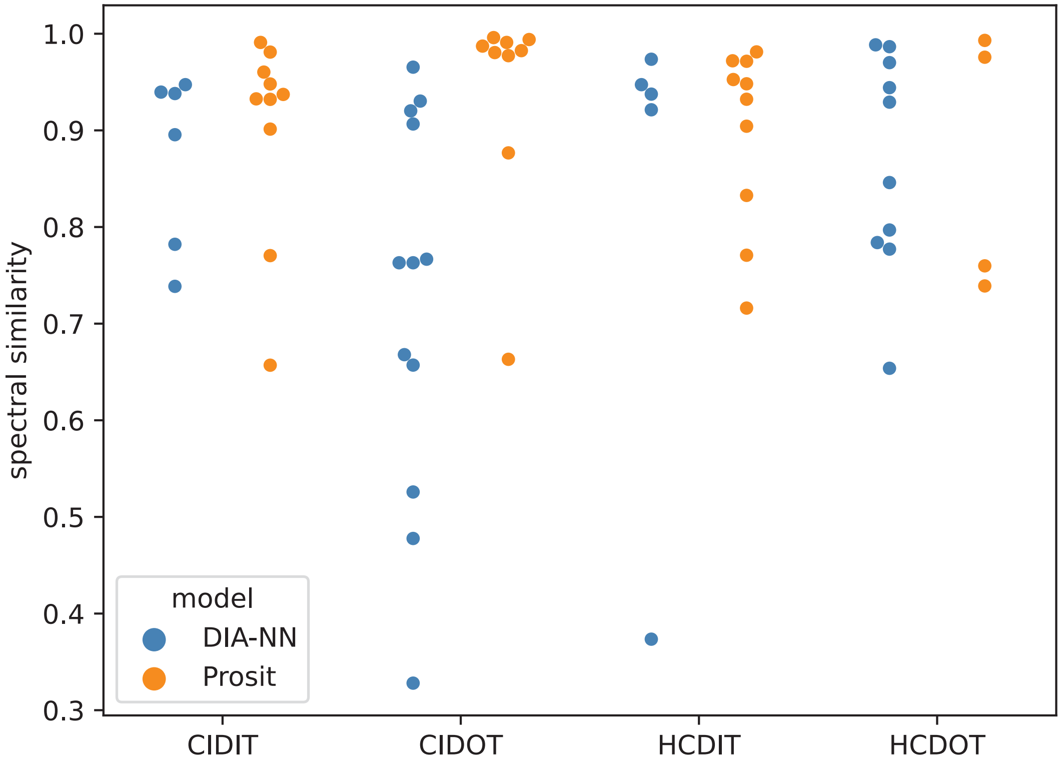


Supplemental Figure 10. For the unique peptides from each model, the highest spectral similarity for PSMs from each peptide are plotted.

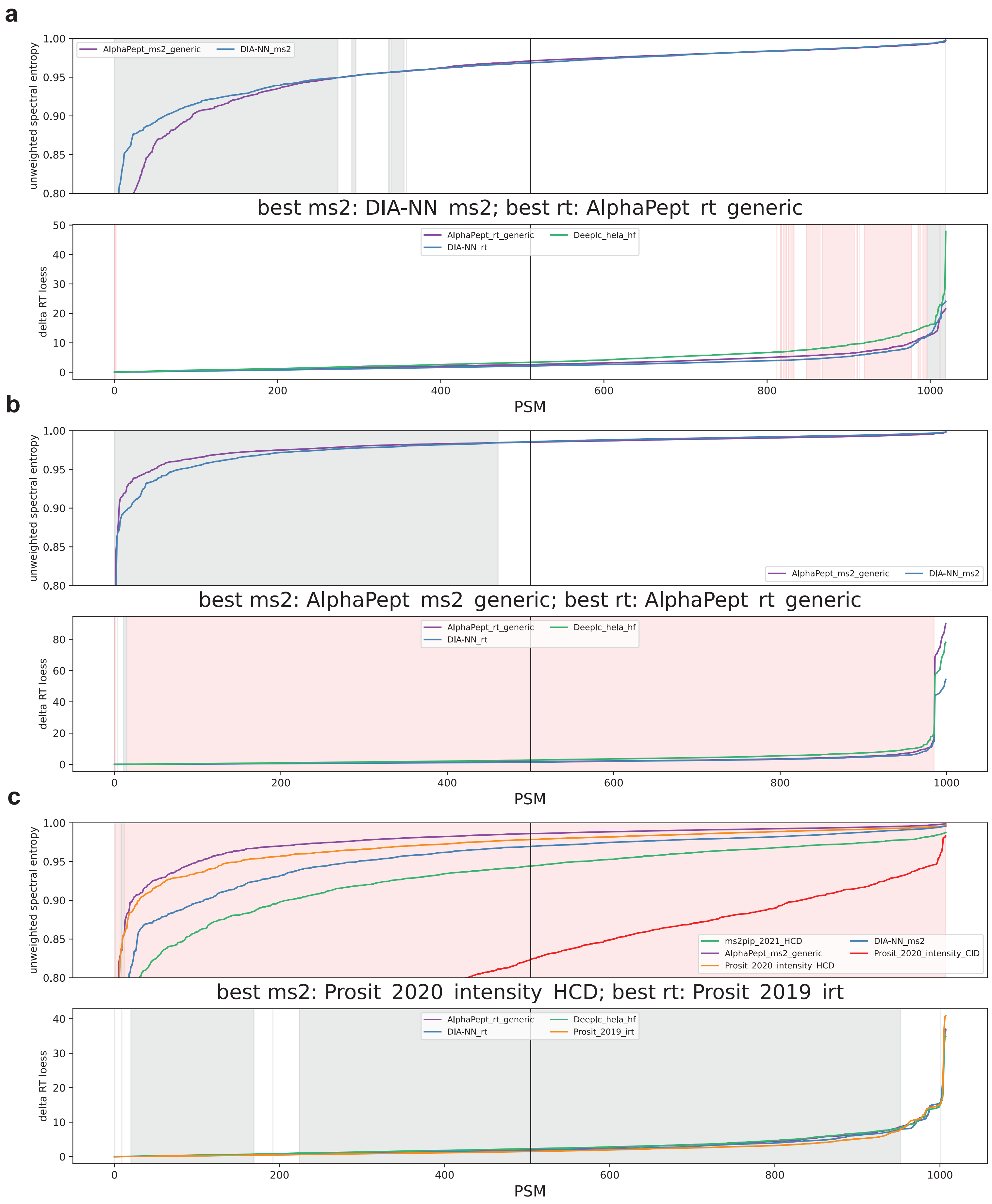

a
unweighted spectral entropy
AlphaPept_ms2_generic
DIA-NN_ms2
best ms2: DIA-NN_ms2; best rt: AlphaPept_rt_generic
delta RT loess
AlphaPept_rt_generic
DIA-NN_rt
Deeplc_hela_hf
PSM
b
unweighted spectral entropy
AlphaPept_ms2_generic
DIA-NN_ms2
best ms2: AlphaPept_ms2_generic; best rt: AlphaPept_rt_generic
delta RT loess
AlphaPept_rt_generic
DIA-NN_rt
Deeplc_hela_hf
PSM
c
unweighted spectral entropy
ms2pip_2021_HCD
AlphaPept_ms2_generic
Prosit_2020_intensity_HCD
DIA-NN_ms2
Prosit_2020_intensity_CID
best ms2: Prosit_2020_intensity_HCD; best rt: Prosit_2019_irt
delta RT loess
AlphaPept_rt_generic
DIA-NN_rt
Deeplc_hela_hf
Prosit_2019_irt
PSM

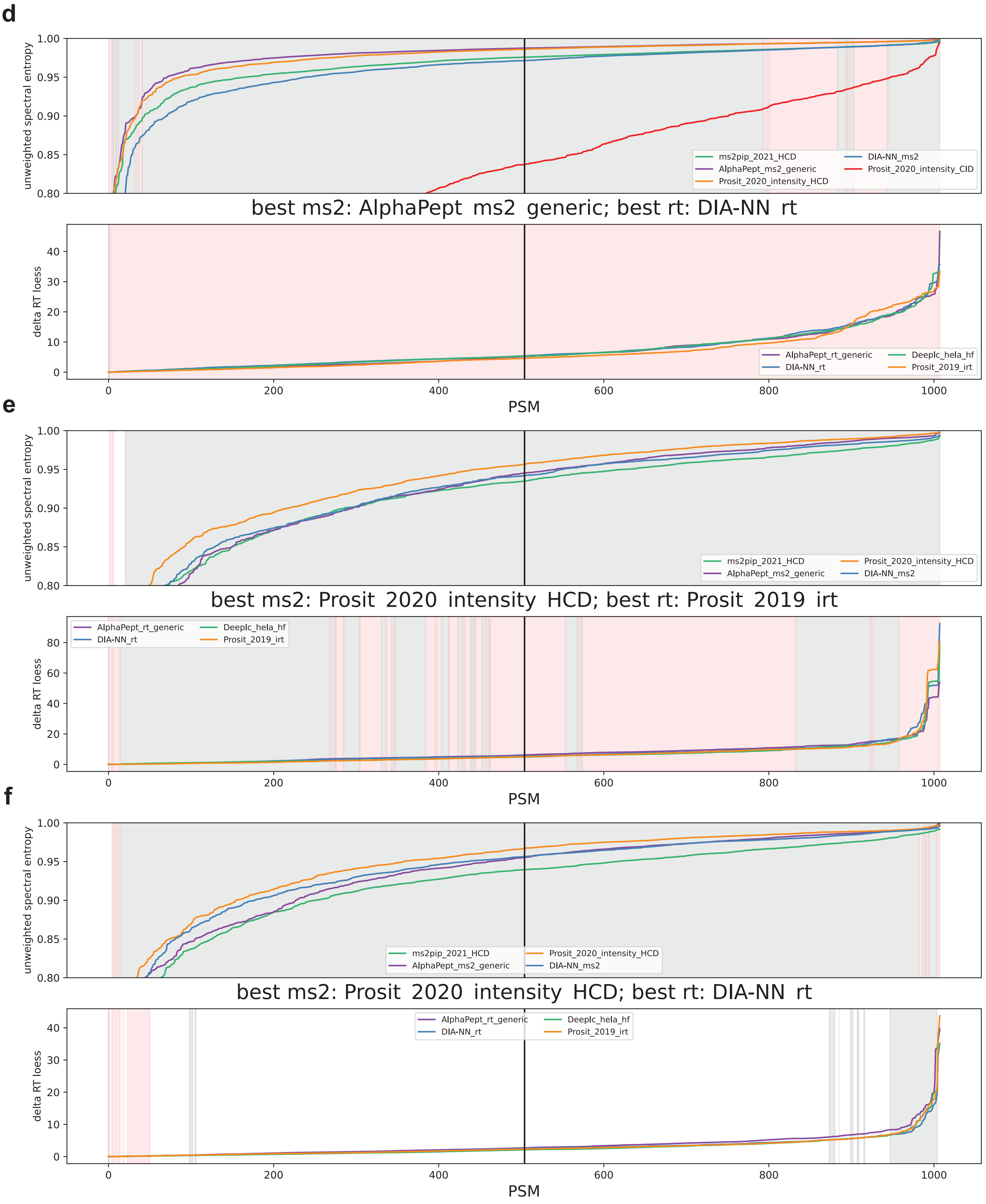
d
unweighted spectral entropy
1.00
0.95
0.90
0.85
0.80
ms2pip_2021_HCD
AlphaPept_ms2_generic
Prosit_2020_intensity_HCD
DIA-NN_ms2
Prosit_2020_intensity_CID
best ms2: AlphaPept_ms2_generic; best rt: DIA-NN_rt
delta RT loess
40
30
20
10
0
AlphaPept_rt_generic
DIA-NN_rt
Deeplc_hela_hf
Prosit_2019_irt
0
200
400
600
800
1000
PSM
e
unweighted spectral entropy
1.00
0.95
0.90
0.85
0.80
ms2pip_2021_HCD
AlphaPept_ms2_generic
Prosit_2020_intensity_HCD
DIA-NN_ms2
best ms2: Prosit_2020_intensity_HCD; best rt: Prosit_2019_irt
delta RT loess
80
60
40
20
0
AlphaPept_rt_generic
DIA-NN_rt
Deeplc_hela_hf
Prosit_2019_irt
0
200
400
600
800
1000
PSM
f
unweighted spectral entropy
1.00
0.95
0.90
0.85
0.80
ms2pip_2021_HCD
AlphaPept_ms2_generic
Prosit_2020_intensity_HCD
DIA-NN_ms2
best ms2: Prosit_2020_intensity_HCD; best rt: DIA-NN_rt
delta RT loess
40
30
20
10
0
AlphaPept_rt_generic
DIA-NN_rt
Deeplc_hela_hf
Prosit_2019_irt
0
200
400
600
800
1000
PSM

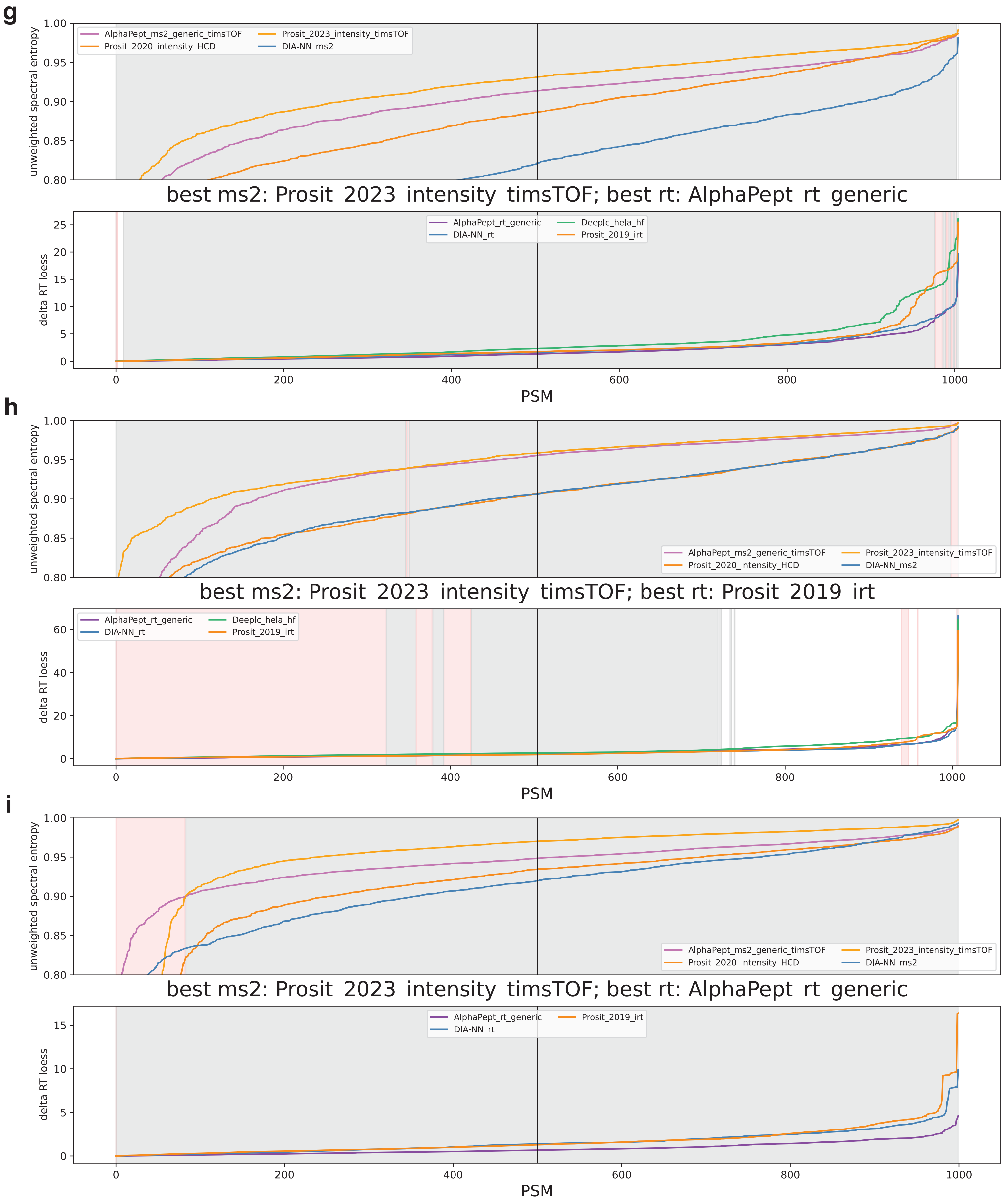
g
unweighted spectral entropy
AlphaPept_ms2_generic_timsTOF
Prosit_2023_intensity_timsTOF
Prosit_2020_intensity_HCD
DIA-NN_ms2
best ms2: Prosit_2023_intensity_timsTOF; best rt: AlphaPept_rt_generic
delta RT loess
AlphaPept_rt_generic
Deeplc_hela_hf
DIA-NN_rt
Prosit_2019_irt
PSM
h
unweighted spectral entropy
AlphaPept_ms2_generic_timsTOF
Prosit_2023_intensity_timsTOF
Prosit_2020_intensity_HCD
DIA-NN_ms2
best ms2: Prosit_2023_intensity_timsTOF; best rt: Prosit_2019_irt
delta RT loess
AlphaPept_rt_generic
Deeplc_hela_hf
DIA-NN_rt
Prosit_2019_irt
PSM
i
unweighted spectral entropy
AlphaPept_ms2_generic_timsTOF
Prosit_2023_intensity_timsTOF
Prosit_2020_intensity_HCD
DIA-NN_ms2
best ms2: Prosit_2023_intensity_timsTOF; best rt: AlphaPept_rt_generic
delta RT loess
AlphaPept_rt_generic
Prosit_2019_irt
DIA-NN_rt
PSM

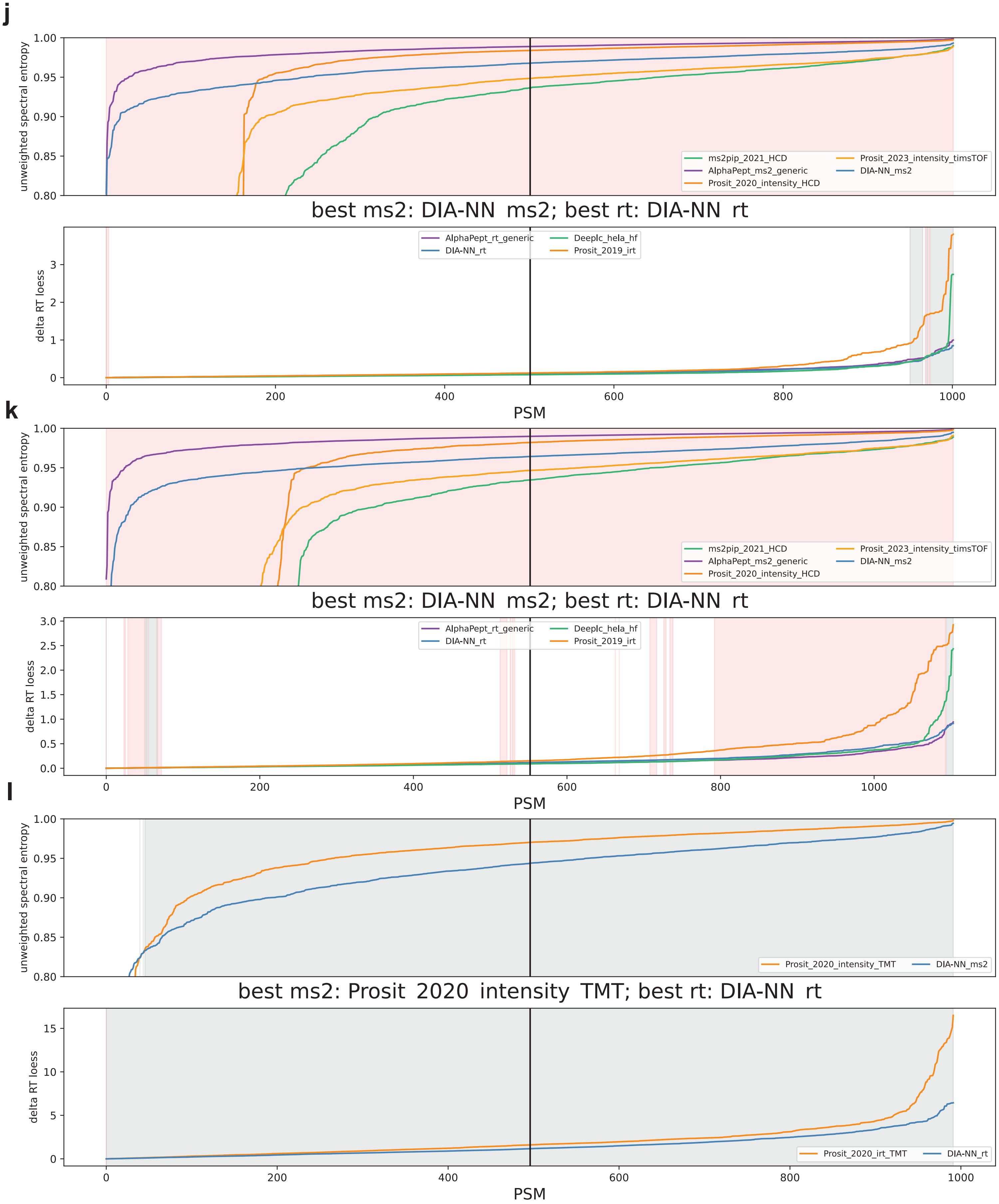


Supplemental Figure 11. Feature values for top PSMs for each model tested during the heuristic model search, just as in Fig 5. The datasets covered here are Arabidopsis phosphoproteome (a), mouse phosproteome (b), HLA DDA class II from Marcu et al., 2021 (c), HLA DDA class I from Pak et al., 2021 (d), HLA DIA class I from Pak et al., 2021 (e) and from Ritz et al., 2017 (f), HLA DDA class I and II on timsTOF SCP (g-h), HeLa tryptic digest on timsTOF (i), Astral DIA from Serrano et al., 2024 (j) and Guzman et al. 2024 (k), and TMT11 LUAD (l).
